# Discovery of small molecule inhibitors of the PTK7/β-catenin interaction targeting the Wnt signaling pathway in colorectal cancer

**DOI:** 10.1101/2021.10.15.464507

**Authors:** Laetitia Ganier, Stéphane Betzi, Carine Derviaux, Philippe Roche, Christophe Muller, Laurent Hoffer, Xavier Morelli, Jean-Paul Borg

**Affiliations:** Aix Marseille Univ, CNRS, INSERM, Institut Paoli-Calmettes, CRCM, Equipe labellisée Ligue ‘Cell polarity, Cell signaling and Cancer’, Marseille, France; Aix Marseille Univ, CNRS, INSERM, Institut Paoli-Calmettes, CRCM, team ‘Integrative Structural and Chemical Biology’, Marseille, France; Aix Marseille Univ, CNRS, INSERM, Institut Paoli-Calmettes, CRCM, ‘HiTS/IPCdd – High throughput screening platform’, Marseille, France; Institut Universitaire de France

**Author notes:** **Corresponding Authors Xavier Morelli** – Aix Marseille Univ, CNRS, INSERM, Institut Paoli-Calmettes, CRCM, team ‘Integrative Structural and Chemical Biology’, 13009 Marseille, France; Aix Marseille Univ, CNRS, INSERM, Institut Paoli-Calmettes, CRCM, ‘HiTS/IPCdd – High throughput screening platform’, 13009 Marseille, France; **Jean-Paul Borg** - Aix Marseille Univ, CNRS, INSERM, Institut Paoli-Calmettes, CRCM, Equipe labellisée Ligue ‘Cell polarity, Cell signaling and Cancer’, 13009 Marseille, France; Institut Universitaire de France. Evotec – 195 Route d’Espagne, 31100 Toulouse, France (Christophe Muller). Drug Discovery Program, Ontario Institute for Cancer Research, Toronto, ON, Canada.

## Abstract

Second cause of death due to cancer worldwide, colorectal cancer (CRC) is a major public health issue. The discovery of new therapeutic targets is thus essential. The pseudokinase PTK7 intervenes in the regulation of the Wnt/β-catenin pathway signaling, in part, through a kinase-domain dependent interaction with the β-catenin protein. PTK7 is overexpressed in CRC; an event associated with metastatic development and reduced survival of non-metastatic patient. In addition, numerous alterations have been identified in CRC inducing constitutive activation of Wnt/β-catenin pathway signaling through β-catenin accumulation. Thus, we thought that targeting PTK7/β-catenin interaction could be of interest for future drug development. In this study, we have developed a NanoBRET™ screening assay recapitulating the interaction between PTK7 and β-catenin to identify compounds able to disrupt this protein-protein interaction. A high-throughput screening allowed us to identify small molecule inhibitors targeting the Wnt pathway signaling and inducing anti-proliferative effect *in vitro* in CRC cells harboring *β-catenin* or *APC* mutations downstream of PTK7. Thus, inhibition of the PTK7/β-catenin interaction could represent, in the future, a new therapeutic strategy to inhibit cell growth dependent on Wnt signaling pathway. Moreover, despite a lack of enzymatic activity of its tyrosine kinase domain, targeting the PTK7 kinase domain-dependent functions appears to be of interest for further therapeutic development.

## INTRODUCTION

Colorectal cancer (CRC) is the second deadliest cancer worldwide with almost 900.000 deaths estimated in 2018 by the GLOBOCAN 2018^1^. Survival of patients with metastatic CRC has significantly improved over the past two decades with an average overall survival of 30 months. This is mainly due to the use of chemotherapies such as oxaliplatin and irinotecan but also to the introduction of targeted therapies. However none of them has shown survival improvement in the adjuvant setting^2,3^. Discovery of new therapeutic targets via a better characterization of molecular key players of CRC is thus essential.

CRC cells frequently reactivate the Wnt pathway involved in the establishment and maintenance of cell polarity, fundamental for the embryonic development of vertebrates and invertebrates. The Wnt pathway is classically divided into two pathways: (i) the canonical pathway also called Wnt/β-catenin and (ii) the non-canonical pathway independent of β-catenin, itself subdivided into two pathways, Wnt/Planar Cell Polarity and Wnt/Ca^2+^ pathways^4^. Reactivation of the Wnt pathway in cancer cells plays an important role at different steps of the tumoral process such as auto-renewal of cancer stem cells, tumorigenesis, metastatic dissemination and resistance to treatments^5^. Alterations of multiple actors of the Wnt signaling, including *Adenomatous Polyposis Coli* (APC), β-catenin or Axin, are found in 93% of CRC^6^. Most of these alterations induce constitutive activation of Wnt/β-catenin pathway signaling through β-catenin accumulation and transcription of target genes involved in tumorigenesis such as *c-MYC*, *CCND1* or *MMP7*^7–9^. Despite increasing attention paid to Wnt/β-catenin pathway in therapeutic development, few molecules have reached clinical trial to date^10^.

The tyrosine kinase receptor PTK7 is a cell surface component of the Wnt pathway^11^. PTK7 was first identified in human normal melanocytes and in colon carcinoma^12,13^. PTK7 is composed of seven extracellular immunoglobulin domains, a transmembrane region and an intracellular tyrosine kinase domain. However, its kinase domain lacks enzymatic activity and therefore this receptor is considered as a pseudokinase^14^. In retrospective studies, PTK7 was found overexpressed in CRC, an event associated with metastatic development, reduced metastases-free survival of non-metastatic patients, and resistance to chemotherapy. Moreover, PTK7 has promigratory and pro-metastatic functions *in vitro* and *in vivo.* However, the mechanisms behind are not well understood yet^15,16^. PTK7 appears to be a promising new therapeutic target. A unique anti-PTK7 therapeutic approach has so far reached the clinical phases and consists in the injection of an antibody-drug conjugate (ADC) targeting tumor cells overexpressing PTK7^17^. Here, PTK7 behaves as a cell surface antigen enabling the destruction of PTK7-positive cancer cells. Development of alternative strategies to ADC through the development of chemical compounds targeting the PTK7 kinase domain-dependent functions would be of great interest. Indeed, we and others have shown that the kinase domain is endowed with signaling functions despite lack of enzymatic activity^18–22^. The role of PTK7 in Wnt/β-catenin pathway has been demonstrated in several studies. Peradziryi *et al* have shown an inhibitory role of PTK7 on the Wnt/βcatenin signaling, which was confirmed in zebrafish^23,24^. However, two other studies including one from our group identified LRP6 and β-catenin as new partners of PTK7 and have correlated loss of PTK7 with inhibition of Wnt/β-catenin signaling^18,25^. We demonstrated that PTK7 binds to β-catenin using a yeast two hybrid assay. Moreover we showed that this direct interaction is mediated by the kinase domain of PTK7 and that it is required for β-catenin-dependent transcriptional events^18^.

In this report, we have developed a NanoBRET™ screening assay which recapitulates the interaction between PTK7 and β-catenin *in cellulo* and is applicable for high throughput screening (HTS) of chemical compounds able to disrupt the PTK7/β-catenin interaction^26,27^. We used different strategies based on virtual screening^28^, HTS of the ‘Fr-PPIChem’ library (10,314 compounds)^29^ and repurposing of TCF/βcatenin inhibitors^30^. These strategies allowed us to identify PTK7/β-catenin small molecules inhibitors targeting Wnt pathway signaling in CRC cells as evidenced in cell-based NanoBRET™ and TopFlash/Luciferase reporter assays. These compounds exhibit an *in vitro* anti-proliferative effect on CRC cells harboring *β-catenin* or *APC* mutations downstream of PTK7, showing the potential of PTK7/ β-catenin targeting for future drug development.

## RESULTS AND DISCUSSION

### Development and validation of a NanoBRET™ assay monitoring PTK7/β-catenin interaction in living cells and optimization for high-throughput screening

In this study, we developed a NanoBRET™ assay to confirm the PTK7/β-catenin interaction and to identify small molecular modulators of the complex in a cellular environment. This technology enables the sensitive and reproducible detection of protein interactions *in cellulo* and is applicable for drug screening^31^. The NanoBRET™ system is a proximity-based assay that can detect protein interactions by measuring energy transfer from a bioluminescent NanoLuc® (NL) fusion protein as an energy donor to a fluorescently labeled HaloTag® (HT) fusion protein as an energy acceptor (**Figure 1A**)^26^. Initial development of the PTK7/β-catenin NanoBRET™ assay was performed with the assistance of Promega Company. Different constructs were generated by appending the HT acceptor tag to PTK7 in N- or C-terminal. The different combinations were tested and PTK7-HT + NL-β-catenin were identified as the optimal pair which yielded an optimal donor to acceptor BRET ratio at 1:10 (**Figure 1B**). Donor Saturation Assay (DSA) showed excellent specificity and assay window. The specific interaction generated a hyperbolic curve where all donor molecules are co-paired with acceptor molecules indicative of a specific BRET interaction detection. Interaction with an unfused HT protein alone as negative control generated a much weaker linear ratio (**Figure 1C**). Following this, an acceptor to donor ratio of 10:1 was used for further assay validation and screening as it lays within the ideal dynamic range for the detection of PTK7/β-catenin interaction changes. To validate the NanoBRET assay for HTS, HEK293 cells transiently transfected with PTK7-HT and NL-β-catenin plasmids, were plated in 384-well plates. The HaloTag 618 ligand was added to generate a positive control signal whereas DMSO alone was added for the negative control signal. The assay statistics evaluation exhibited a 0.69 Z’-factor value and a >2 signal to background ratio (**Figure 1D**). Taken together, these values confirmed that we were able to develop a specific and sensitive PTK7/β-catenin NanoBRET™ interaction assay compatible with high-throughput screening applications.

**Figure 1.**
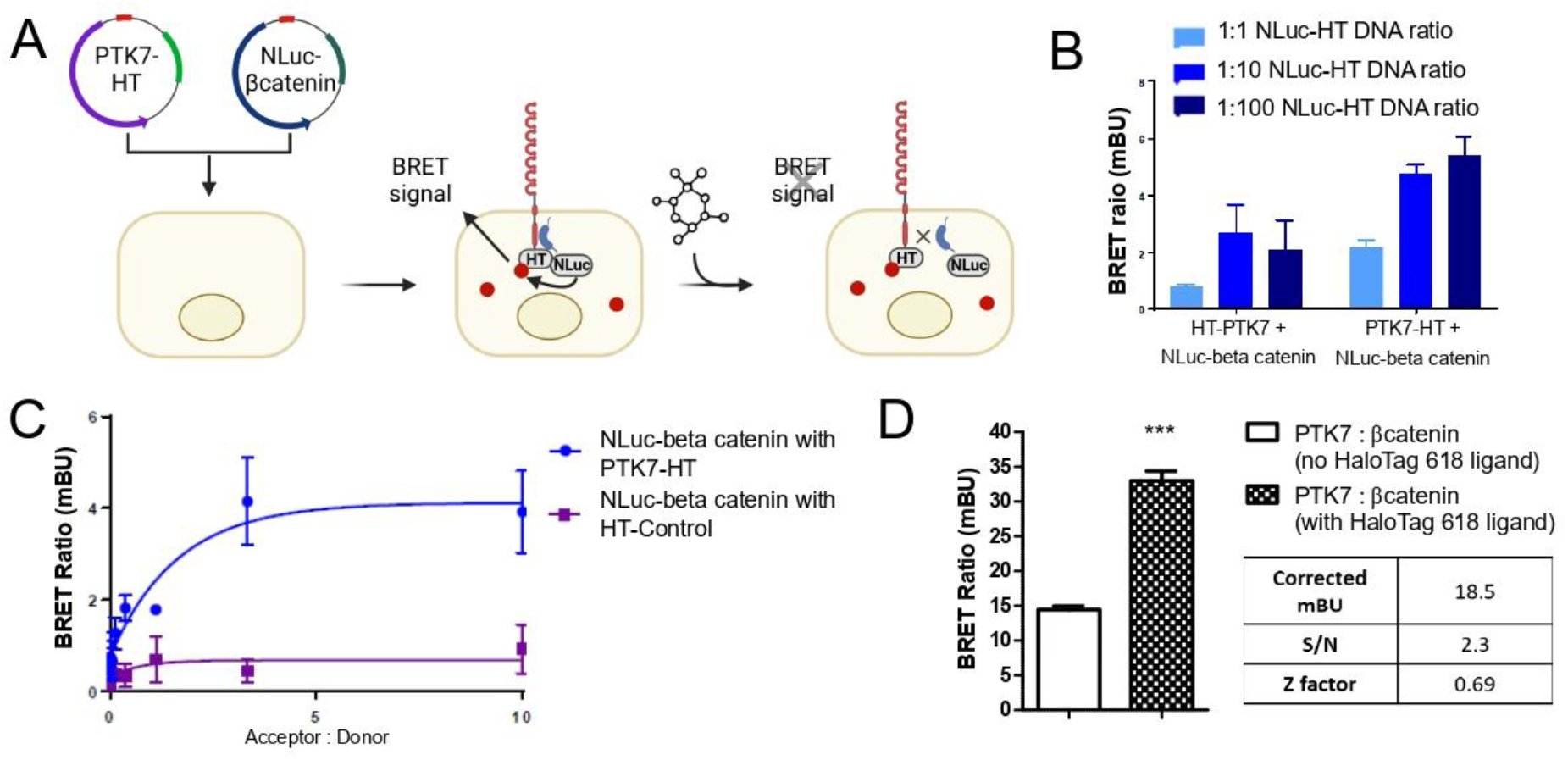
Development and validation of a NanoBRET assay monitoring PTK7/β-catenin interaction in living cells and optimization for high-throughput screening. (A) Scheme illustrating the proximity-based NanoBRET assay. PTK7-HT (HaloTag) and NL (NanoLuc)-β-catenin expressing plasmids are transfected in HEK293T cells. Cells are then incubated with HaloTag 618 ligand (red circle) and Furimazine (substrate of the Nano-Luciferase) is added to allow the generation of a BRET signal. Molecules that disrupt the PTK7/β-catenin complex yield reduced BRET signal. (B) N- and C-terminal HT fusions to PTK7 with NL-β-catenin were tested at three donor to acceptor ratios. (C) Donor Saturation Assay (DSA) showing specificity for the detection of the interaction between NL-β-catenin and PTK7-HT. Cells were transfected with constant amount of NL donor DNA and paired with increasing amounts of acceptor DNA to obtain increasing amounts of Acceptor-to-Donor (A/D) ratios (blue line). The negative control is an unfused HaloTag® protein used as mock acceptor (purple line). (D) Optimization of the NanoBRET PTK7/β-catenin interaction assay for HTS. HEK293T cells expressing PTK7-HaloTag and NanoLuc-β-catenin were plated in a 384-well plate, at a density of 8,000 cells per well, with HaloTag 618 ligand to generate a positive BRET signal or with DMSO as negative control. Data represents 176 (positive control) and 16 (negative control) wells, with mean ± SD (t test, ***p<0.001). Donor-to-acceptor ratios in positive and negative controls were used to calculate the Signal-to-Noise ratio (S/N). The Z-factor is defined with means (μ) and standard deviations (σ) of both positive (p) and negative (n) controls. The Z-factor is defined as: Z-factor = 1 – [3(σ_p_ + σ_n_)/(μ_p_ – μ_n_)]. A S/N ratio >2. A Z-factor value between 0.5 and 1 is suitable for HTS.

### Use of different approaches to identify small-molecule inhibitors of the PTK7/β-catenin interaction

To identify chemical compounds that inhibit the PTK7/β-catenin protein-protein interaction (PPI), we developed a multi-screening strategy based on our newly developed NanoBRET assay that combined virtual screening, HTS and repurposing of drug in development as summarized in **Figure 2A**.

**Figure 2.**
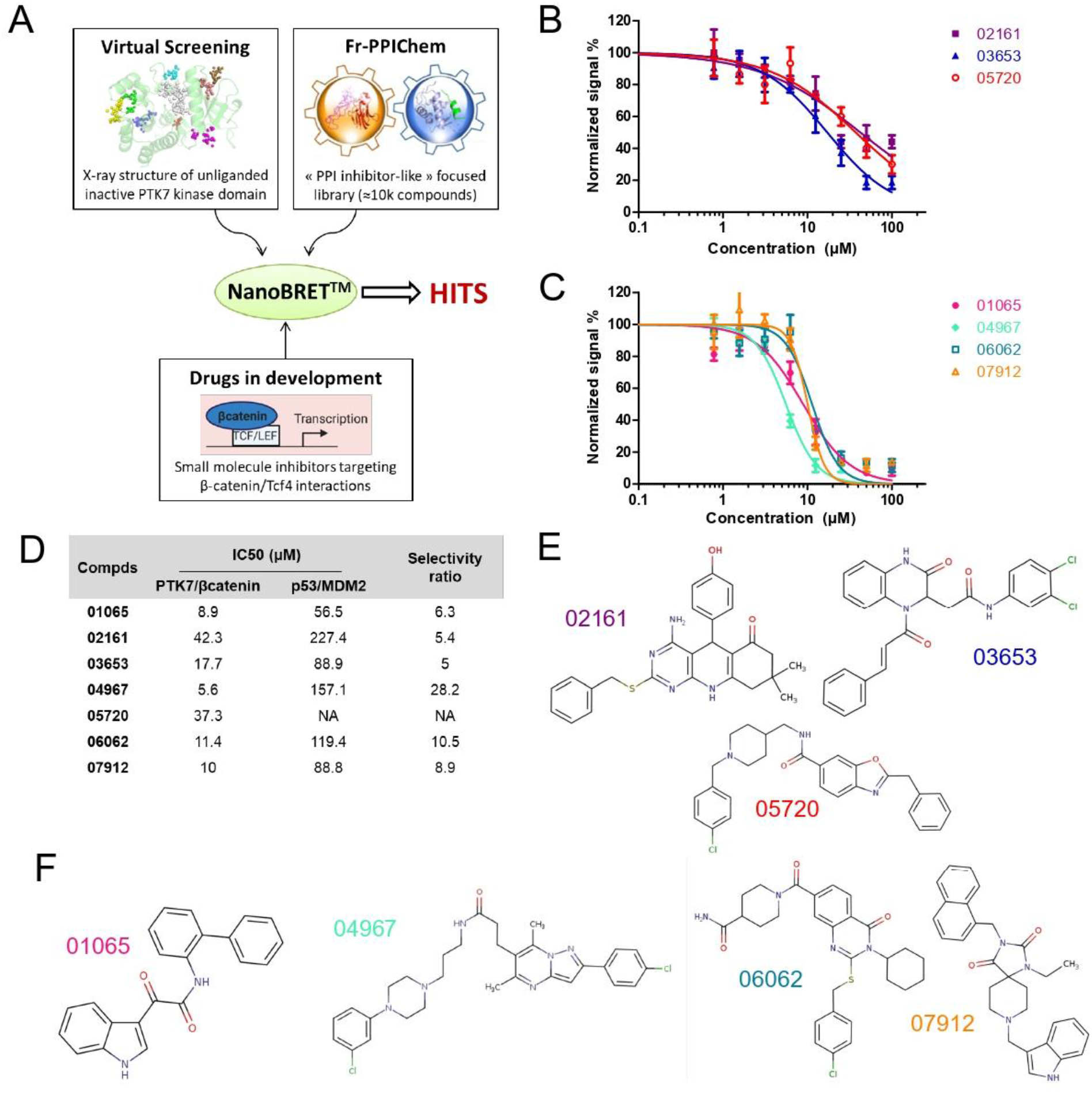
Use of different approaches to identify small-molecule inhibitors of the PTK7/β-catenin interaction. (A) Illustration of the different strategies used to identify hits: virtual screening, random screening and repurposing of drugs in development. (B-C) BRET signal in transiently transfected HEK293 cells with PTK7-HT and NL-β-catenin plasmids incubated overnight with increasing concentrations of compounds. Values are mean ± SD, n=3. NanoBRET IC_50_ assays allow the identification of 3 hits from the virtual screening strategy (D) and 4 hits from the Fr-PPIChem HTS. (D) Inhibitor selectivity of small molecule PTK7/β-catenin inhibitors over p53/MDM2 interaction determined by NanoBRET™. (E-F) Structures of PTK7/βcatenin inhibitors identify from the virtual screening (E) and the random screening (F).

In a first approach, we took advantage of the recently published X-ray structure of unliganded inactive PTK7 kinase domain to perform an *in silico* screening of the FrPPIChem library (10,314 compounds). This library has been designed to contain potential protein-protein interaction inhibitor (iPPI) compounds^14,29^. This structure was used as a starting template to identify potential direct binders of PTK7. The experimental approach was based on a strategy described previously^28^ and combines three robust computational approaches: Molecular Dynamics (MD) simulations, molecular docking methods, and pharmacophore filtering. The analysis of the MD trajectory of PTK7 protein identified conformations with putative larger druggable pockets that might be able to interact with small molecule compounds (**Figure S1A**). Then, high-throughput molecular docking of the Fr-PPIChem library using PLANTS^32^ and MOE (https://www.chemcomp.com/) led to the selection of the 200 best ligands (consensus scoring strategy). Finally, pharmacophore filtering using LigandScout^33^, led to the selection of 92 prioritized compounds that were purchased and tested (**Figure S1B**).

Second, we performed an unbiased screening of the FrPPIChem library that could identify also β-catenin binders with a risk of loss of specificity. Indeed, β-catenin is the target of small molecule ligands^30^ and contains central armadillo repeats responsible for binding with most of its partners including the transcription factors Tcf/Lef, APC and E-cadherin^30^.

Third, we hypothesized that small molecule inhibitors of the β-catenin/Tcf4 interaction already in development could also block the PTK7/β-catenin interaction and serve as a basis for future optimization of the specificity. We selected some β-catenin/Tcf4 inhibitors with different specificity profiles towards Tcf4 (**Table S1**).

In the end, a total of 10,413 compounds selected from these three complementary strategies were purchased and evaluated using our optimized PTK7/β-catenin NanoBRET assay. From this HTS campaign, 3 compounds were identified as inhibitor hits from the *in silico* approach with IC_50_ values ranging from 15 to 40μM (**Figure 2B**) and 4 additional compounds from the random screening of the Fr-PPIChem library with IC_50_ values ranging from 5 to 10μM (**Figure 2C**). Interestingly, we could not identify compounds among the β-catenin/TCF4 inhibitors having inhibitory effect on the PTK7/β-catenin interaction even for less selective compounds (**Table S1**). This observation suggests a different binding site of PTK7 to β-catenin compared to Tcf4, APC or E-cadherin.

In addition, we confirmed hit compounds selectivity by performing a NanoBRET™ assay on another optimized control pair of vectors designed to measure the interaction between p53 and MDM2 (**Figure S2**). All compounds exhibit significant selectivity for PTK7/β-catenin over p53/MDM2 interaction (>5-fold). Compound 04967 exhibited the best inhibitory potential on the PTK7/β-catenin interaction with a 5.6 μM IC_50_, as well as the highest selectivity ratio versus p53/MDM2 interaction with a 28.2 value (**Figure 2D**). Overall, the NanoBRET™ HTS campaign allowed the identification of seven hits (**Figure E-F**) inhibiting the PTK7/β-catenin interaction with micromolar range potency and good selectivity versus an irrelevant complex.

### SAR-by-catalog resulting in the identification of PTK7/β-catenin inhibitors targeting Wnt pathway signaling in CRC cells

In order to overcome potential false positive and identify compounds with lower IC_50_, we performed a structure-activity relationship study by purchasing commercially available compounds related to our hits (SAR-by-catalog). For this, we ordered 5 to 13 analogs for each of our 7 compounds and NanoBRET™ IC_50_ assays were performed for all purchased analogs. Only 2 series of derivatives exhibited clear structure activity relationships (modulations of the compound scaffold affect the IC_50_) confirming a specific inhibition of the complex (refer to **Table S2** for more details regarding the 2D chemical structures and IC_50_ information for all analogs). The first series includes 01065 identified from the Fr-PPIChem library and two of its analogs: 20269 and 20274 (**Figure 3A**). The second includes 03653 identified from the *in silico* screening on PTK7 X-ray structure and two of its analogs: 20278 and 20279 (**Figure 3B**). These results prompted us to further investigate how these molecules could modulate Wnt pathway signaling and confirm their potential as molecular probes of the PTK7/β-catenin interaction in cells.

**Figure 3.**
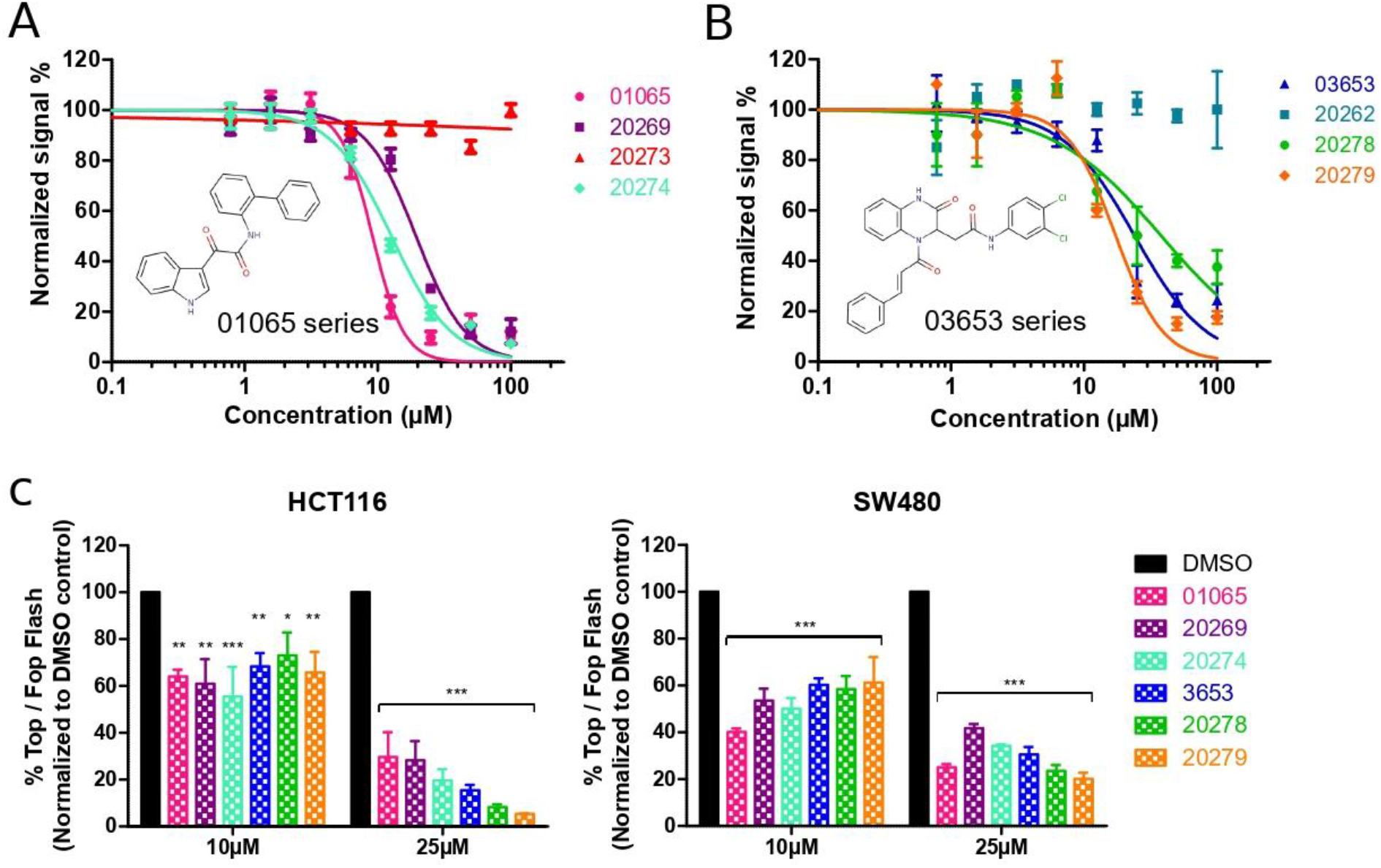
Use of SAR-by-catalog resulted in the identification of PTK7/β-catenin inhibitors targeting the Wnt signaling pathway in CRC cells. (A-B) BRET signal in transiently transfected HEK293 cells incubated overnight with increasing concentrations of hit analogs. Values are mean ± SD, n=3. NanoBRET IC_50_ assays on the 55 analogs allow identification of active analogs only for the 01065 (A) and 03653 (B) hits. (C) Effect of the different compounds on WNT signaling pathway. The CRC cells lines HCT116 and SW480 were incubated overnight with the different compounds at 10 and 25μM. Wnt signaling activity was measured using the TopFlash/Luciferase reporter assay normalized with FOP activity and expressed as percentage of control cells treated with DMSO. Results are expressed as mean of 3 independent experiments ± SD and *P* values are derived from two-way Anova (*p < 0.05; **p <0.01; ***p < 0.001).

As mentioned previously, the role of PTK7 in the Wnt/β-catenin signaling pathway is cell context dependent^18,23–25^. To evaluate PTK7 involvement in the Wnt/β-catenin signaling in CRC, we used two colorectal cancer cell lines (HCT116 and SW480) with different mutational status. HCT116 cells are heterozygous for *β-catenin* with a mutant allele bearing a mutation of a serine involved in the negative regulation of β-catenin following phosphorylation by GSK3^34^. SW480 cells are *APC^mt^* resulting in a C-terminal truncated protein lacking the β-catenin binding domain^35^. Both mutations induce a constitutive activation of the Wnt/β-catenin pathway signaling in HCT116 and SW480. Firstly, these two CRC cell lines were transfected with a potent siRNA against PTK7 (**Figure S3A**) and effect on the Wnt/β-catenin signaling was evaluated using a β-catenin bioluminescent reporter assay (TOPflash normalized with FOPflash). We observed a significant inhibition in the transactivating activity of β-catenin following PTK7 downregulation in the two cell lines (**Figure S3B**). These results are in line with those of Puppo *et al.*^18^ and suggest an inhibitory role of PTK7 on Wnt/β-catenin signaling independently of *β-catenin* or *APC* mutations.

We thus moved to the biological validation of our two series of compounds. CRC cells were treated overnight with all compounds at 10 and 25μM and activity on Wnt signaling was evaluated with the β-catenin bioluminescent reporter assay. All compounds significantly decreased the transactivating activity of β-catenin in HCT116 and SW480 in a dose-dependent manner (**Figure 3C**). As part of the biological validation, an analog from the 03653 series not selected in NanoBRET™ (compound 20262) had no effect on Wnt signaling activation in both cell lines at 25μM (**Figure S3C**). Considering the above results, we conclude that downregulation of PTK7 or PTK7/β-catenin inhibition with small molecules attenuate Wnt signaling activation in CRC cells.

### PTK7/β-catenin inhibition with small molecules modulates Wnt signaling target genes expression in CRC cells

To analyze the effect of our compounds on Wnt signaling pathway, we used a RT^2^ profiler PCR array to analyze the expression of 84-Wnt related target genes following treatment of CRC cells. These 84 genes are listed in **Table S3** with their main associated functions as for example cell adhesion, cell migration, cell cycle regulation, differentiation and development or signal transduction. HCT116 and SW480 cells were treated for 24h with the two hits, 01065 and 03653, at 25μM for gene expression analysis. Analogs were not tested in the preliminary screening based on the assumption that they should have a similar mechanism than their related compound. Results are presented on scatter plots (**Figure 4A-B**); the central line indicates no change in gene expression compared to DMSO control. Gene expression changes are beyond the boundary lines with a fold-change threshold of 2 and are represented in red dots for up-regulated genes or green dots for down-regulated genes. The RT^2^ screening led to the identification of Wnt target genes differentially regulated in a same way in all conditions such as *AXIN2* but also genes modulated in a cell line- or compound-dependent fashion as for *MMP7* downregulated only in HCT116 or *DKK1* upregulated in both cell lines following treatment with compound 01065 only.

**Figure 4.**
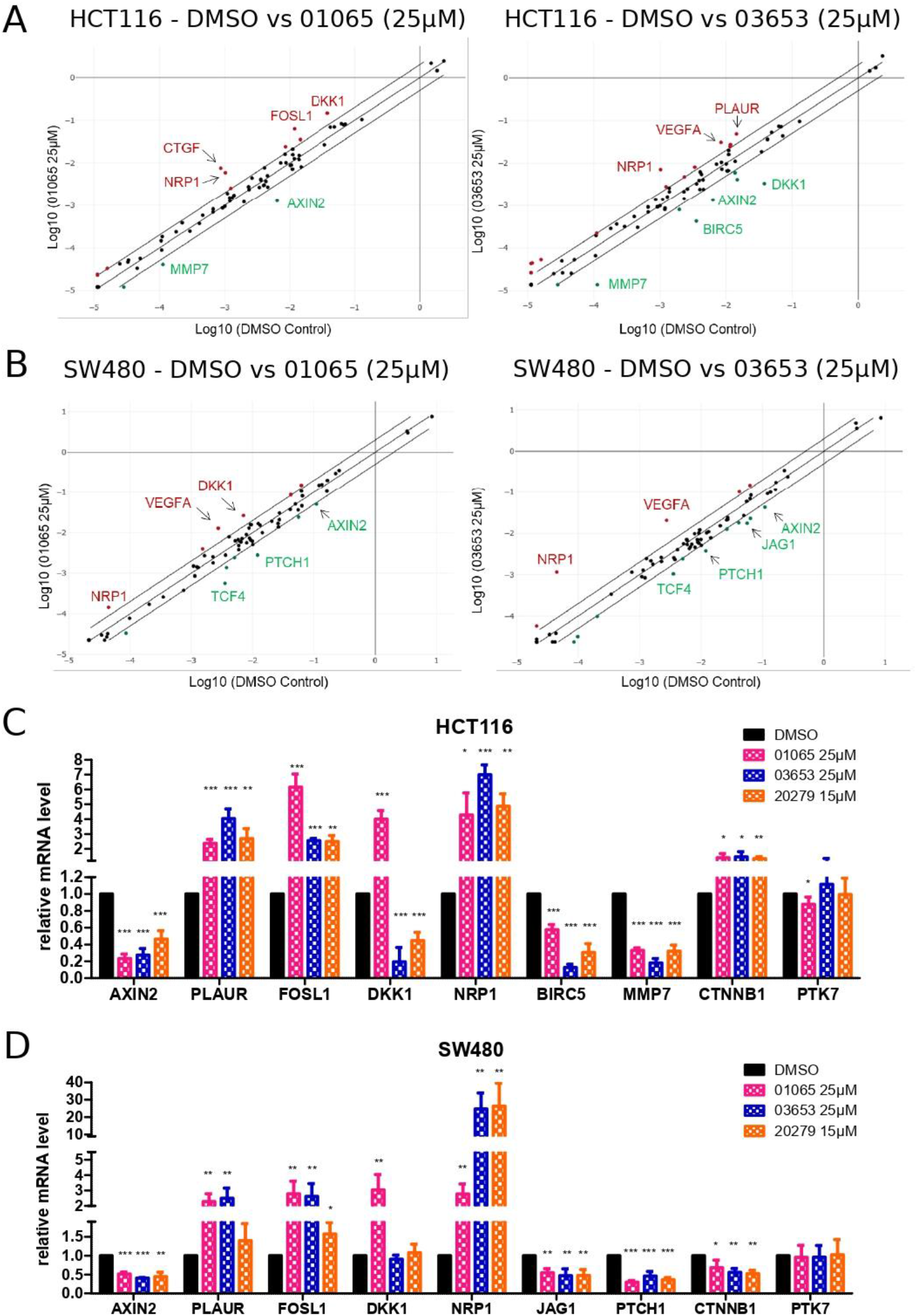
PTK7/β-catenin inhibitors modulate Wnt signaling target genes expression in CRC cells. (A-B) Effects of compounds 01065 and 03653 on Wnt signaling target genes expression profiles in HCT116 (A) and SW480 (B) cells. Cells were treated 24h with compounds 01065 and 03653 at 25μM. The expression change of the 84 genes in samples and control groups was revealed by real-time PCR based array analysis. Scatter plots graph the log_10_ of the expression level of each gene in the sample group (01065 or 03653) versus the corresponding value in the control group (DMSO). The central line indicates fold changes of 1, or no change. The external lines indicate a fold-change threshold of 2. The red and green dots stand for up-regulated and downregulated genes respectively. (C-D) Validation of selected candidate genes with classical RT-qPCR in HCT116 (C) and SW480 (D). Cells were treated 24h with compounds 01065 and 03653 at 25μM. Validation analysis identified compound 20279 as more efficient than its parent hit 03653 and thus treatment was performed at a reduced concentration of 15μM. Other analogs identified in Figure 2 did not show improvement in effect and are not represented in this figure. Data show relative mRNA levels of candidate genes in sample groups against DMSO control group. Results are expressed as mean of 3 independent experiments ± SD and *P* values are derived from t-tests (*p < 0.05; **p <0.01; ***p < 0.001).

We then validated selected genes candidate from gene profiling analysis with classical RT-qPCR in HCT116 (**Figure 4C**) and SW480 (**Figure 4D**) cells. During the validation study, only the 20279 analog appeared to be more efficient than its parent compound and exhibited similar effects at 15μM to its parent compound (03653) at 25μM. We could confirm the significant reduction in *AXIN2* mRNA level in HCT116 and SW480 treated with 01065, 03653 and 20279. AXIN2 is a well-characterized target of misregulated β-catenin responsive transcription that acts as a feedback inhibitor to impede the activity of Wnt signaling pathway^36^. The three compounds also induce a significant upregulation in *PLAUR*, *FOSL1* and *NRP1* mRNA levels in both cell lines. Interestingly, a previous study has shown that siRNA-mediated silencing of β-catenin increased PLAUR expression at the mRNA and protein levels in SW480 cells which is consistent with our results^37^. NRP1 is strongly upregulated especially in SW480 treated with 03653 and 20279 with a fold change > 20. Interestingly, NRP1 was shown to have differential pro-tumoral *vs.* anti-tumoral effects depending on the type of cancer and the mutational status of *KRAS*. It was shown that inhibition of NRP1 expression in cells containing dominant active *KRAS*^mt^ caused increased cell viability and tumor growth^38^. As HCT116 and SW480 are both *KRAS*^mt^, NRP1 upregulation led by 01065, 03653 and 20279 could theoretically reduce tumor growth. The compound 01065 only induces upregulation of *DDK1* mRNA encoding a secreted inhibitor of Wnt signaling, in both cell lines. Other chemical compounds, such as HDAC inhibitors, have been shown to upregulate DKK1 in colon cancer initiating cells and lead to cell cycle arrest and apoptosis^39^. Finally, other genes are specifically modulated in one cell line in the same way with all compounds tested. This includes down-regulation of *BIRC5* (survivin) and *MMP7* in HCT116, and *JAG1* and *PTCH1* in SW480. All these genes play crucial roles in cancer progression and particularly cell proliferation and cell survival so their down-regulation should promote anti-tumorigenic functions. Apart from Wnt target genes, no major effect of the compounds was evidenced on *PTK7* mRNA expression. However, *β-catenin* mRNA levels were down-regulated in SW480 cells. In this cell line, β-catenin cannot be degraded due to APC mutation and, in regards with Wnt/β-catenin signaling inhibition following compounds treatment, *β-catenin* mRNA down-regulation could be part of a transcriptomic negative feedback to reduce β-catenin protein level available in cells. In HCT116, *β-catenin* mRNA levels were slightly up-regulated but are not biologically significant as fold-changes are <2.

Altogether, these data show that PTK7/β-catenin inhibitors induce differential expression of Wnt signaling target genes associated with cell proliferation, cell survival and migration.

### PTK7/β-catenin inhibitors induce differential cytotoxicity between CRC cell lines and Mouse Embryonic Fibroblasts (MEFs)

To investigate the antiproliferative properties of 01065, 03653 and 20279 *in cellulo*, we performed Alamar Blue assays on HCT116 and SW480 treated cells for 72h with increasing concentrations of compounds. All tested compounds exerted dose-dependent antiproliferative effects against CRC cells, compound 20279 being the most active in accordance with previous results (**Figure 5A – Table1**). We also evaluated the specificity over CRC cells by performing this assay on primary Mouse Embryonic Fibroblasts (MEFs) expressing PTK7. At a dose inducing maximum antiproliferative properties on CRC cells (30μM for 01065 and 03653 and 15μM for 20279); weak effect was observed on MEFs (**Figure 5A – Table1**).

**Table 1.** Table recapitulating IC_50_ values (concentration of compounds exhibiting 50% cell viability) in HCT116, SW480 and MEFs.

| IC<sub>50</sub> (μM) | 01065 | 03653 | 20279 |
| --- | --- | --- | --- |
| HCT116 | 19.6 ± 0.59 | 20.1 ± 1.7 | 11.3 ± 0.25 |
| SW480 | 21.3 ± 0.84 | 19.6 ± 0.73 | 10.7 ± 0.30 |
| MEF | 41.0 ± 4.95 | 46.3 ± 4.77 | 21.2 ± 1.32 |

**Figure 5.**
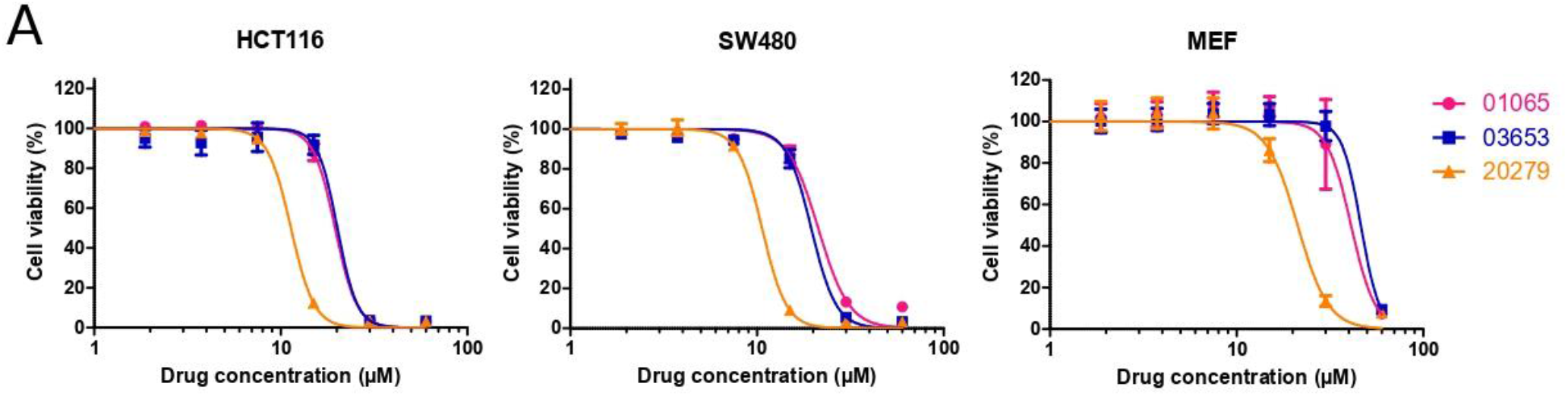
PTK7/β-catenin inhibitors induce differential cytotoxicity between CRC cell lines and primary Mouse Embryonic Fibroblasts (MEFs). (A) Impact of compounds 01065, 03653 and 20279 on HCT116, SW480 and MEFs viability *in vitro*. Cells were incubated for 72h with increasing concentration of compounds. Cell viability was assessed by Alamar Blue assay and expressed as percentage of control cells treated with DMSO. Points represent mean of three independent experiment and bars are SD.

Altogether, PTK7/β-catenin inhibitors demonstrate anti-proliferative activity with highest selectivity over CRC cells. This can be explained by PTK7 overexpression and constitutive activation of the Wnt/β-catenin pathway signaling in CRC cells compared to MEFs which require stimulation by Wnt ligands (for example Wnt3a) to activate the Wnt pathway.

### PTK7/β-catenin inhibitors cause cell cycle arrest of CRC cells

Wnt/β-catenin signaling is well known to promote cell proliferation through transcriptional up-regulation of target genes involved in cell cycle progression. Another evidence of a relation between the cell cycle and Wnt signaling comes from the observations that β-catenin protein levels and Wnt/β-catenin signaling oscillate during the cell cycle^40,41^. Consistently with our previous findings, we thus investigated deeper the antiproliferative effect of our compounds. We performed a cell cycle distribution analysis based on a standard flow cytometry method by staining DNA with propidium iodide (PI) 24h after treatment of HCT116 and SW480 cells. Interestingly, both 01065 and 03653 series induced an obvious cell cycle arrest through distinct mechanisms. On the one hand, compound 01065 increased the percentage of HCT116 and SW480 cells in S phase with subsequent decrease in G0/G1 phases suggesting S phase arrest that inhibits cell cycle progression (**Figure 6A**). On the other hand, the 03653 series strongly increased the percentage of HCT116 cells in G0/G1 phases, more moderately in the case of SW480 cells, suggesting also an inhibition of cell cycle progression (**Figure 6A**).

**Figure 6.**
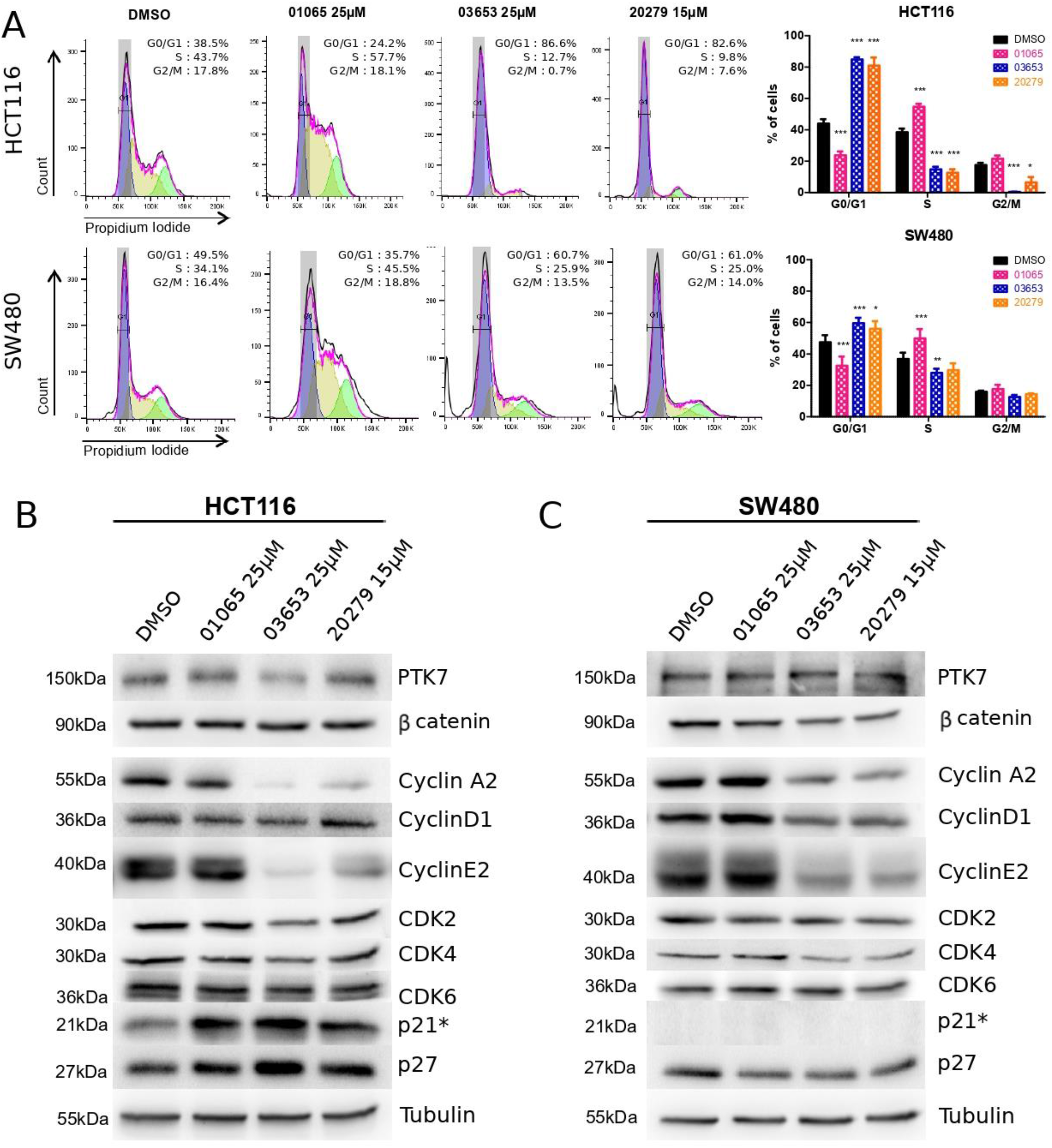
PTK7/β-catenin inhibitors cause cell cycle arrest in CRC cells. (A) Left: Cell cycle distribution of HCT116 and SW480 measured by propidium iodide (PI) staining and analyzed for DNA content by flow cytometry after treatment with DMSO, 25μM of 01065 and 03653 or 15μM of 20279 for 24hours. Right panel: Cell cycle distribution is represented as histogram of the percentage of cells in G0/G1, S or G2/M phase. Results are expressed as mean of 3 independent experiments ± SD and *P* values are derived from two-way Anova (*p < 0.05; **p <0.01; ***p < 0.001). (B-C) Total cell lysates were extracted from HCT116 (B) and SW480 (C) cells to examine the expression levels of PTK7, βcatenin and key proteins involved in cell cycle regulation by western analysis. *To note p53 regulates p21 expression but while HCT116 are p53^wt^, SW480 are p53^mt^ rendering p53 inactive which explains why p21 is not expressed in this cell line^42,43^.

We then determined the levels of key cell cycle regulators in HCT116 and SW480 cells after drug treatment. In HCT116 cells, we could detect a significant increase in p21 and p27 protein levels for both series of compounds. The 03653 series also induced a strong decrease in CyclinA2 and CyclinE2 and a slight decrease in CDK proteins consistent with the observed cell cycle arrest (**Figures 6B and S4A**). Regarding SW480 cells, the results are less clear for the compound 01065 as no major regulation of cell cycle proteins was observed. However, we could detect a significant decrease in CyclinA2, CyclinD1, CyclinE2 and CDK4 but also in p27 with the 03653 series (**Figure 6C and S4B**). Of note, in contrast to HCT116 cells, SW480 cells do not express p21 due to inactive p53 which normally regulates p21^42,43^.

Apart from cell cycle regulators, we could not detect any obvious alteration of PTK7 protein levels in both cell lines. Regarding β-catenin, the compounds did not reduce its protein level in HCT116 cells. One possible explanation for β-catenin responsive transcription inhibition could be an inhibition of β-catenin translocation into the nucleus *via* formation of E-cadherin/β-catenin complexes, for example, as it was shown in another study^44^. In SW480 cells, β-catenin protein levels are slightly decreased and only significantly upon treatment with compound 03653 (**Figures 6B-C**). However, this decrease could be due to *β-catenin* mRNA downregulation (**Figure 4C**).

Compound 01065 induced cell cycle arrest at S phase whereas 03653 and 20279 compounds induced cell cycle arrest at G0/G1 phases and consequently inhibited HCT116 and SW480 cell proliferation. However, the HCT116 cell line appears to be more sensitive to PTK7/β-catenin inhibitors than the SW480 cell line. One hypothesis could be that the β-catenin/PTK7 expression ratio is around 1 in SW480 while it is 0.5 in HCT116 (data not shown). Thus, an increase of PTK7/β-catenin complexes in SW480 could reduce compounds efficiency.

We also assessed if these compounds could induce cell apoptosis following cell cycle arrest. We performed a standard flow cytometry analysis by staining cells with Annexin V and propidium iodide (PI) 48 hours after treatment. However, we could not observe induction of cell apoptosis in the two cell lines suggesting that effect of our compounds is restricted to cell proliferation (**Figures S5A-B**). Association of these compounds with proapoptotic compounds could be of interest as concomitant growth arrest and apoptosis are required for effective targeted therapy^45,46^.

In conclusion, we demonstrated that inhibition of the PTK7/β-catenin interaction is achievable with small molecules and could represent, in the future, a new therapeutic strategy to inhibit CRC cell growth dependent on Wnt signaling pathway. Additional studies will be required, particularly with regard to the potency of these compounds. These compounds also represent interesting molecular probes that can be used to target and modulate PTK7/βcatenin interaction and better understand this interaction in its pathways and in cancer development.

## Supporting information

Supporting Information Ganier et al

## ASSOCIATED CONTENT

### Supporting Information.

The Supporting Information is available free of charge via the internet at http://pubs.acs.org.

Cell lines and reagents; experimental methods for *in silico* structure-based screening, NanoBRET™ assay, TopFlash assay, cell viability assay, RT^2^ profiler PCR arrays, flow cytometry analysis and immunoblot analysis; *In silico* screening workflow, hit compounds selectivity, role of PTK7 on Wnt pathway signaling, western blot quantification, cell apoptosis analysis; List of 7 small molecule inhibitors targeting β-catenin/TCF4 interaction, List of 55 compounds to probe the SAR of hit compounds, List of 84 key genes responsive to Wnt signal transduction, primer sequences and antibodies.

## Author Contributions

LG and SB designed the project, performed experiments, analyzed data and wrote the manuscript. CD manages the High Throughput Screening platform HiTS/IPCdd and helped for the HTS screening. CM performed the virtual screening. LH and PR performed the SAR-by-catalog analysis. JPB and XM supervised, analyzed, funded the project and wrote the manuscript. All authors have given approval to the final version of the manuscript.

## Notes

The authors declare no potential conflicts of interest.

## ACKNOWLEDGEMENT

The authors wish to thank the R&D department from Promega for the generation of the NanoBRET™ plasmids and validation of the assay. We also thank La ligue Nationale Contre le Cancer and Fondation pour la Recherche Médicale for their support.

---

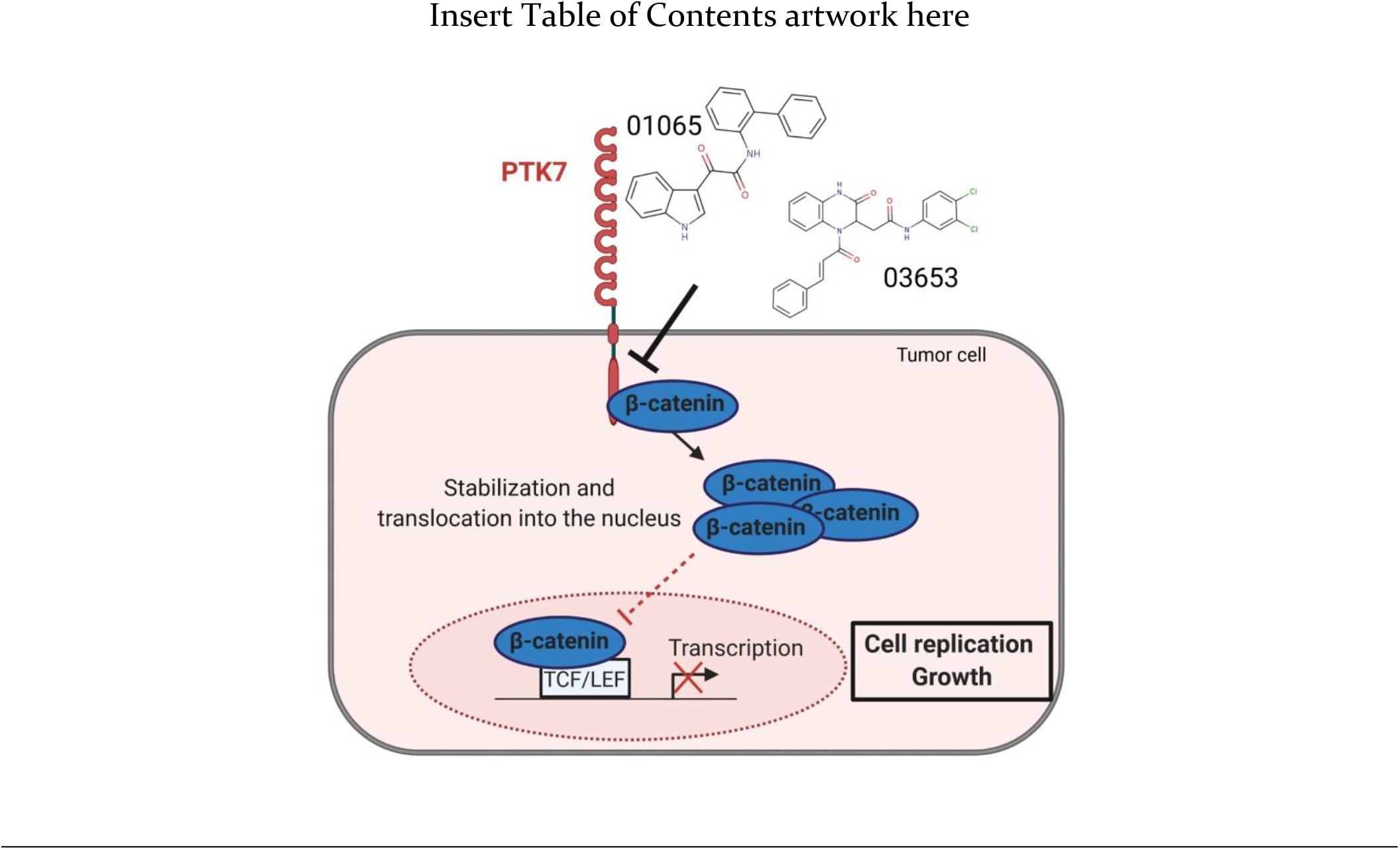

