## Supporting Information Ganier et al for "Discovery of small molecule inhibitors of the PTK7/β-catenin interaction targeting the Wnt signaling pathway in colorectal cancer"

- **Table S1**: Small molecule inhibitors targeting β-catenin/TCF4 interactions tested in NanoBRET for their cross-reactivity on the PTK7/β-catenin interaction 12

### I. MATERIALS AND METHODS

#### Cell lines and reagents

HEK293T and SW480 were cultured in DMEM supplemented with 10%FBS. HCT116 were cultured in RPMI supplemented with 10% FBS. Mouse embryo fibroblasts (MEFs) were cultured in DMEM supplemented with 15% FBS, 1% sodium pyruvate, 1% non-essential amino acids and 50µM 2-mercaptoethanol. All cells were grown at 37°C under 5% $CO_2$ with relative humidity.

The Fr-PPIChem library resulted from a French consortium including present authors to create a unique chemical library dedicated to the inhibition of protein-protein interactions (PPIs)[1]. The Fr-PPIChem library is available upon request in 384-well plates.

The siRNA sequence for PTK7 used was no. 25 : 5'-ACGUGGUAGUAGCGAGGUA-3' (Horizon Discovery Ltd).

#### In silico structure-based screening

The *in-silico* screening was designed based on a strategy described previously and combines three robust computational approaches : molecular dynamics (MD) simulations, molecular docking and pharmacophore filtering [2]. We performed MD simulations of PTK7 in periodic conditions in explicit aqueous solvent using the CHARMM force field starting from the X-ray structure of unliganded, inactive PTK7 kinase domain (pdb code : 6VG3)[3]. The initial structure was prepared with MOE software (https://www.chemcomp.com/). The protein was solvated in a TIP3P water box containing chloride and sodium ions to ensure a neutral system. This step and inputs for MD production stage were prepared with the Charmm-gui server (https://www.charmm-gui.org/). Electrostatic interactions were treated using the particle-mesh Ewald summation method, and we used the switch function for the van der Waals energy interactions with cuton, cutoff and cutnb values of 10, 12 and 14 Å, respectively. The Shake algorithm was applied to all hydrogen-containing bonds. The system was equilibrated at 330K for 2 ns followed by a production phase of 50ns using 2 fs integration step. Coordinates were saved every 0.5 ps for subsequent analysis.

We performed a clustering analysis of the MD ensemble to reduce the number of conformations to be used as receptors in high-throughput docking. The clustering analysis was mainly aimed at separating the 'open' from 'closed' conformations and thus revealing cryptic pockets. KMeans was used as the clustering method and a threshold similarity of 1.5Å was chosen. Analysis of cavities presence and frequency of opening among all frames with F-pocket[4] and MDpocket[5] led to the identification of two druggable pockets more accessible based on the RMSD. The high-throughput molecular docking step was performed on the surface of the previously 10 minimized and optimal PTK7 protein structures using PLANTS[6] and MOE (https://www.chemcomp.com/) on the 10,314 compounds from the Fr-PPIChem library. A maximum of 10 poses were generated by the docking algorithm for each compound followed by selection of the top ranked pose for each compound. PLANTS initial scoring and MOE rescoring led to a consensus and selection of the 200 best ligands. Each generated model was then subjected to a structure-based pharmacophore filtering using LigandScout[7]. Criteria for compound selection were more than 2 interactions or original interaction within the protein cavity leading to a final set of 100 compounds of which 92 were available in the MolPort database (https://www.molport.com/). These compounds were purchased and tested experimentally by NanoBRET™.

**NanoBRET™ assay**

The initial development of the NanoBRET™ assay to detect the interaction of PTK7 and β-catenin in transiently-transfected HEK293 cells was done with Promega. The NanoBRET™ protocol was adapted from a technical manual optimized by Promega for the detection of PTK7/βcatenin interaction.

HEK293T cells were resuspend to a final density of $4x10^5$ cells/mL in cell culture medium and 2mL (800,000 cells) were plated into a well of a six-well plate. Cells were allowed to attach and recover for 4-6 hours at 37°C, 5% CO2 and humidity. Cells were transfected using Lipofectamine® LTX & PLUS™ reagent (ThermoFischer Scientific). A first transfection mixture containing 300µL of Opti-MEM® Reduced Serum Medium (no phenol red) with 2µg of PTK7 acceptor vector, 0,2µg of β-catenin donor vector and 3µL of PLUS™ reagent and a second one containing 300µL of Opti-MEM® Reduced Serum Medium phenol-free with 6µL of Lipofectamine® LTX reagent were prepared. The mixtures are mixed and incubated at room temperature for 10 minutes. The transfection mixture is added to wells with attached cells and proteins are expressed for approximately 20-24hours at 37°C, 5% CO2 and humidity. For larger scale experiments, cells were transfected in $10cm^2$ dishes with adapted quantity of cells and transfection reagents accordingly.

The next day, cells were trypsinized, washed in complete media, and re-suspended in OptiMEM phenol-free containing 4% FBS at a density of $2x10^5$ cells/mL. HaloTag 618 ligand was then added to the cell suspension (1µL/mL of 0.1mM stock solution), and 39,5µL of the cell mixture was plated per well of a 384-well white-walled tissue culture plate. For the "no-acceptor controls", 1μL of DMSO were added per mL of cells. When needed, compounds are added in 384-well white-walled tissue culture plate beforehand at the desired concentration with the Labcyte Echo® Acoustic Liquid Handling and adjusted to a final concentration of 1%DMSO (maximum tolerated). For vehicle control, 1% DMSO was added to the cells. Plates were then incubated overnight for maximum compound effect. To measure BRET signal, the Nano-Glo substrate was diluted in OptiMEM phenol-free (10μL/mL) and 10μL of the mixture was added to each well. Donor (460nm) and acceptor (618nm) emissions were measured using the PHERAstar® FSX microplate reader (BMG LABTECH). To calculate raw NanoBRET™ ratio values, the acceptor emission value is divided by the donor emission value for each sample.

**TopFlash assay**

Canonical Wnt/β-catenin signaling activation was evaluated using transient transfection of reporter plasmids. HCT116 ($1 \times 10^5$ cells/well) and SW480 ($7 \times 10^4$ cells/well) were seeded in 24-well plates and transfected with siPTK7 #25 using Lipofectamine® RNAi max (ThermoFischer Scientific) when needed. After 48h, cells were transfected with 500ng of TK-luciferase (firefly) reporter plasmid containing wild-type (TOPflash) TCF/LEF binding sites or containing mutated TCF/LEF binding sites (FOPFlash) and 70ng of Renilla luciferase reporter plasmid (as transfection efficiency reporter) using Lipofectamine® LTX and plus reagent (ThermoFischer Scientific). These plasmids were a nice gift from Christophe Ginestier. 8h later transfection medium was replaced wish fresh medium containing the different drugs at 10 and 25μM or 1% DMSO for incubation overnight. Renilla and Firefly luciferase expression were revealed using Dual-Luciferase Reporter assay (Promega) according to the manufacturer's instructions and measured using a CLARIOstar microplate reader (BMG LABTECH). Data are expressed as (TOPflash Luciferase Firefly/Renilla ratio) / (FOPflash Luciferase Firefly/Renilla ratio) ratio normalized to DMSO treated cells.

**Cell viability assay**

HCT116 ($7 \times 10^3$ cells/well), SW480 ($5 \times 10^3$ cells/well) and MEFs ($1 \times 10^4$ cells/well) were seeded in 96-well plates (Corning®) and incubated for 24h. Cells were treated with a range of concentrations of drugs with a dilution factor of 2 (1,87-60 μM). After 72 hours drug incubation, metabolic activity was detected by addition of 25μL/well of Alamar Blue and fluorescence intensity was measured using a CLARIOstar microplate reader (BMG LABTECH). Cell viability was determined and expressed as a percentage of control cells treated with 1% DMSO.

**RT-qPCR**

HCT116 and SW480 cells were seeded in 6-well plates (Corning®) and incubated for 24h. Cells were treated with candidate inhibitors or DMSO for 24h. Total RNA was isolated using the RNeasy mini kit (Qiagen) according to the manufacturer's instructions and treated with DNase1. Total RNA was quantified on NanoDROP spectrophotometer ND-1000 (Thermo Fischer Scientific). After heat activation, total RNA was reverse transcribed to the first-strand cDNA using SuperScript™ II First-Strand Synthesis system (Thermo Fischer Scientific) with random primers (Thermo Fischer Scientific). The reverse transcribed product is used to run real time PCR reactions using SYBR Green mastermix (ThermoFischer Scientific) on a Biorad CFX96™ real-time cycler. Primers obtained from Life Technologies/ThermoFischer Scientific are listed in **Table S4**.

**RT$^2$ Profiler PCR arrays**

Preparation of samples and RNA extraction are done similarly than in RT-qPCR section. Single-stranded cDNA is synthesized from all samples using the RT$^2$ First Strand Kit (Qiagen) as described in the Qiagen protocol for RT$^2$ profiler array sample preparation. The reverse transcribed product is used to run real time PCR reactions using RT$^2$ SYBR Green qPCR mastermix (Qiagen) on a Biorad CFX96™ real-time cyclers.

**Flow cytometry analysis**

HCT116 and SW480 cells were seeded in 6-well plates (Corning®) and incubated for 24h. For analysis of the cell cycle, cells were treated with candidate inhibitors or DMSO for 24h. After treatment, cells were trypsinized and permeabilize with the eBioscience™ Foxp3 / Transcription Factor Fixation / Permeabilization kit (ThermoFisher Scientific). Briefly, cells were incubated for 20min at 4°C in 1mL of permeabilization buffer. Then, cells were washed in 2mL of Wash buffer 1X diluted in PBS-1% BSA. Cells were stained for 20min at RT in 200µL of wash buffer 1X containing 40µg/mL of propidium iodide and 40µg/mL of RNase A (DNase and protease-free) (ThermoFisher Scientific) and analyzed on a BD™ LSRII flow cytometer.

For apoptosis analysis, cells were treated with candidate inhibitors or DMSO for 48h. Cells were stained for 15min at RT in 100µL of Annexin-binding buffer containing 1µg/mL of propidium iodide and 5µL/assay of Annexin V FITC and analyzed on a BD™ LSRII flow cytometer.

**Immunoblot analysis**

Cells were lysed in ice-cold lysis buffer containing 50mM Tris pH7.5, 0,5M NaCl, 1% Igepal, 1% Deoxycholic acid, 0,1% SDS, 2mM EDTA, Proteinase and Phosphatase Inhibitor Cocktail. Typically, 25µg cell lysates were migrated in SDS-PAGE (Sodium Dodecyl Sulfate-PolyAcrylamide Gel Electrophoresis), with variable % depending on the proteins analyzed, and transferred onto Amersham™ Protran™ Supported 0.45 µM or 0.2µM NC (GE Healthcare BioSciences). The primary antibodies used are listed in **Table S5**.

**Statistical analysis**

All graphs and statistical analyses were done using the GraphPad Prism™ software. All data are presented as mean of multiple experiments (+/- standard deviation). Statistical tests used for determination of P-values are specified in corresponding figure legends. For all analyses, only P-values <0.05 were considered as statistically significant, the key for asterisk placeholders for P-values in the figures are: ***P<0.001, ** P<0.01, * P<0.05.

### II. SUPPLEMENTARY FIGURES

**Figure S1** (Ganier *et al.*)

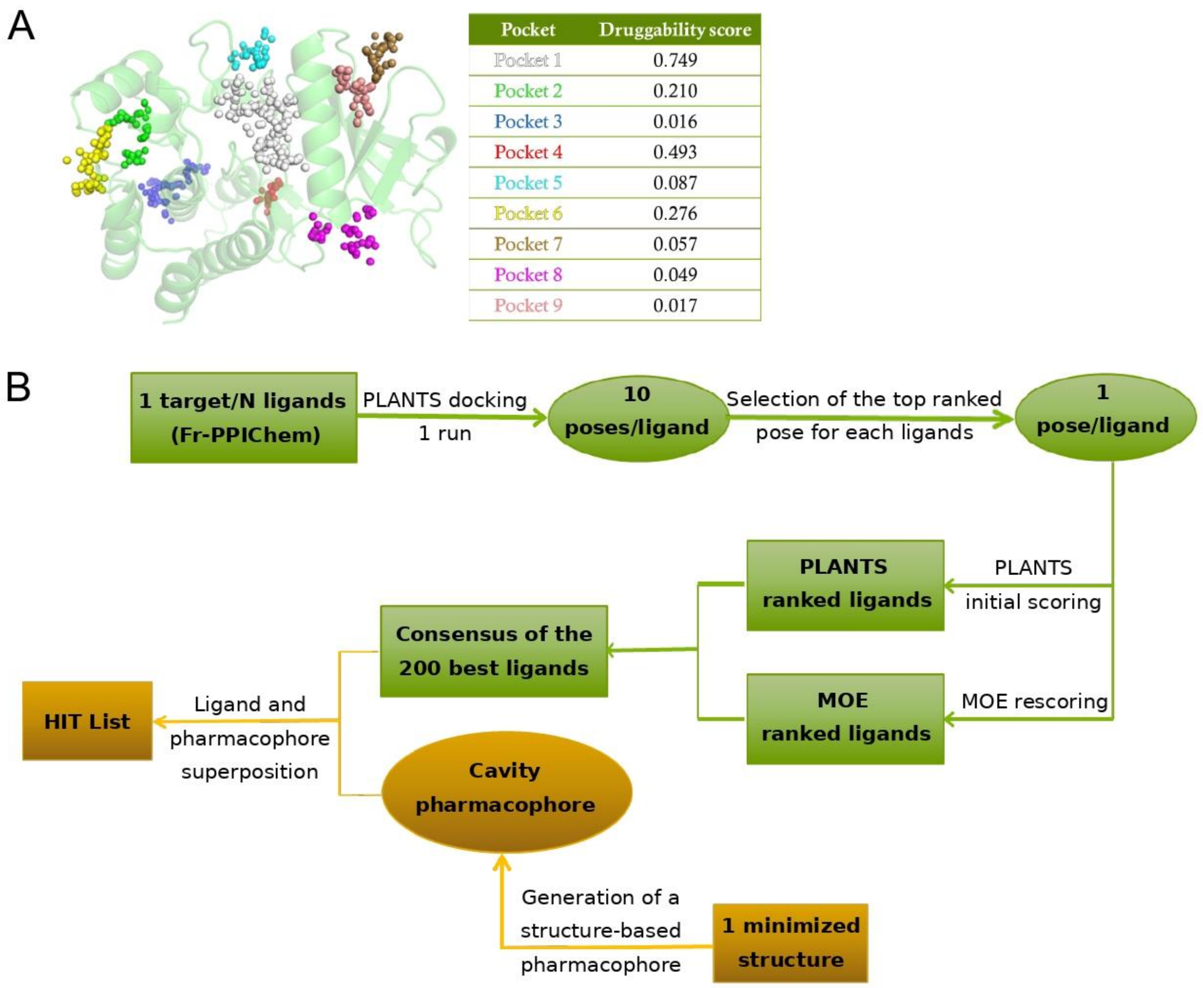

**Figure S1**. (A-B) ***In silico* screening workflow** (A) The first step consisted in MD simulations of PTK7 to account for the structural flexibility of the protein and reveal potential cryptic pockets. Pockets detected in the MD conformations starting from the X-ray structure of unliganded inactive PTK7 kinase domain with FPocket/MDPocket are shown together with the associated druggability score. Pockets 1 and 2 were selected for subsequent docking studies based on the accessibility of these pockets to small molecules. (B) The second step consisted in high-throughput molecular docking of the Fr-PPIChem library[1]. This step was performed on the surface of optimal PTK7 protein structures identified previously using PLANTS[6] and MOE (https://www.chemcomp.com) to generate several compounds conformations that would fit each compound of the chemical library into the two binding pockets identified. This led to a consensus and selection of the 200 best ligands. Finally, the last step subjected each generated model to our structure-based pharmacophore filtering using LigandScout[7] which led to our final hit list.

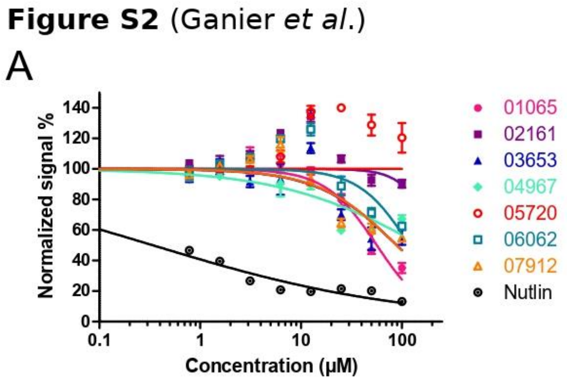

**Figure S2. Evaluation of hit compounds selectivity** (A) BRET signal in transiently transfected HEK293 cells with p53-HT and NL-MDM2 plasmids incubated overnight with increasing concentrations of compounds. Values are mean ± SD, n=3.

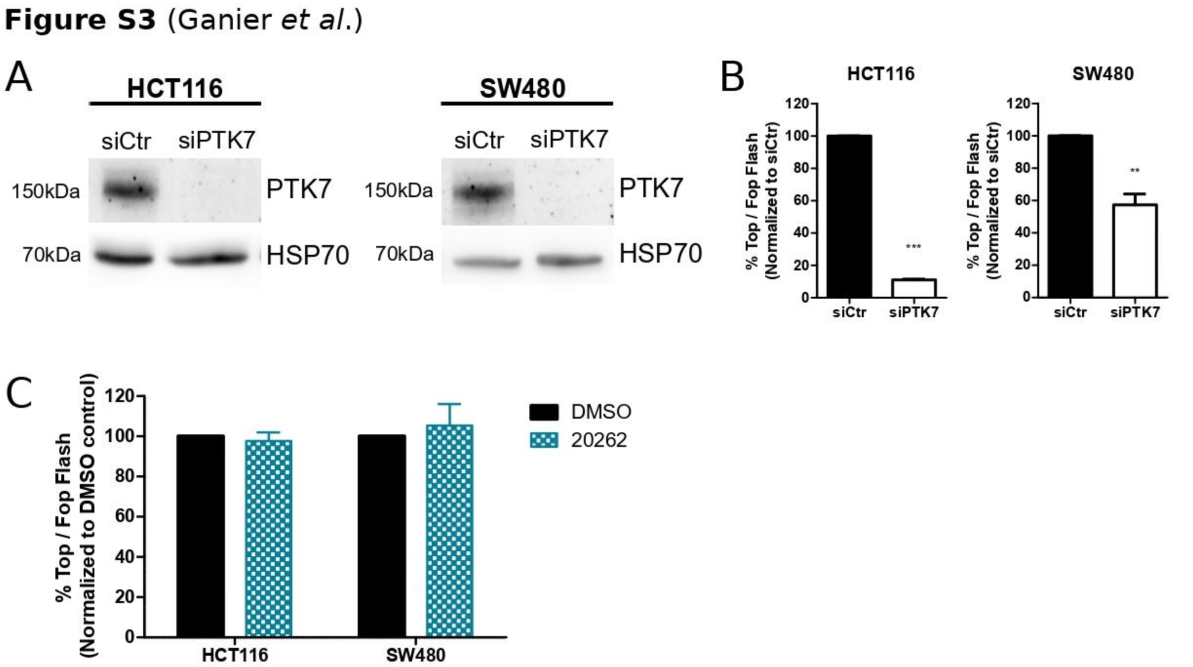

**Figure S3. Identification of PTK7/β-catenin inhibitors targeting Wnt pathway signaling in CRC cells** (A) PTK7 siRNA KD efficiency was evaluated by western blot analysis. HSP70 was used as a loading control. (B) WNT signaling measured by TopFlash assay normalized with FOP activity on HCT116 and SW480 cells treated for 48h with siPTK7 or siCtr. (C) HCT116 and SW480 were incubated overnight with the compound 20262 at 25µM. WNT signaling activity was measured as reported in Figure 2C.

**Figure S4** (Ganier *et al.*)

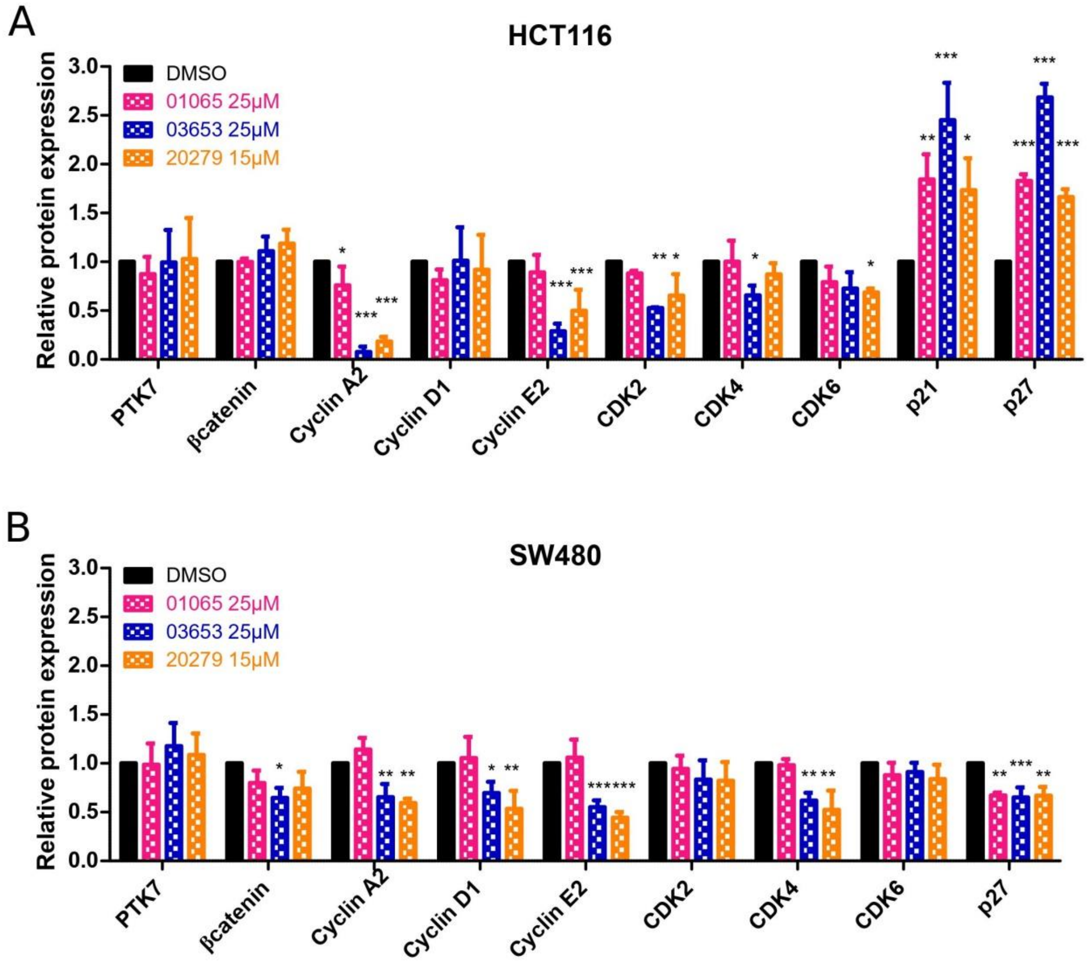

**Figure S4. PTK7/β-catenin inhibitors cause cell cycle arrest in CRC cells.** (A-B) Quantifications of western blot analysis in HCT116 (A) and SW480 (B) were performed using Image J software and are represented as histograms of relative protein expression normalized to DMSO control. Results are expressed as mean of 3 independent experiments ± SD and *P* values are derived from one-way Anova (*p < 0.05; **p <0.01; ***p < 0.001).

**Figure S5** (Ganier *et al.*)

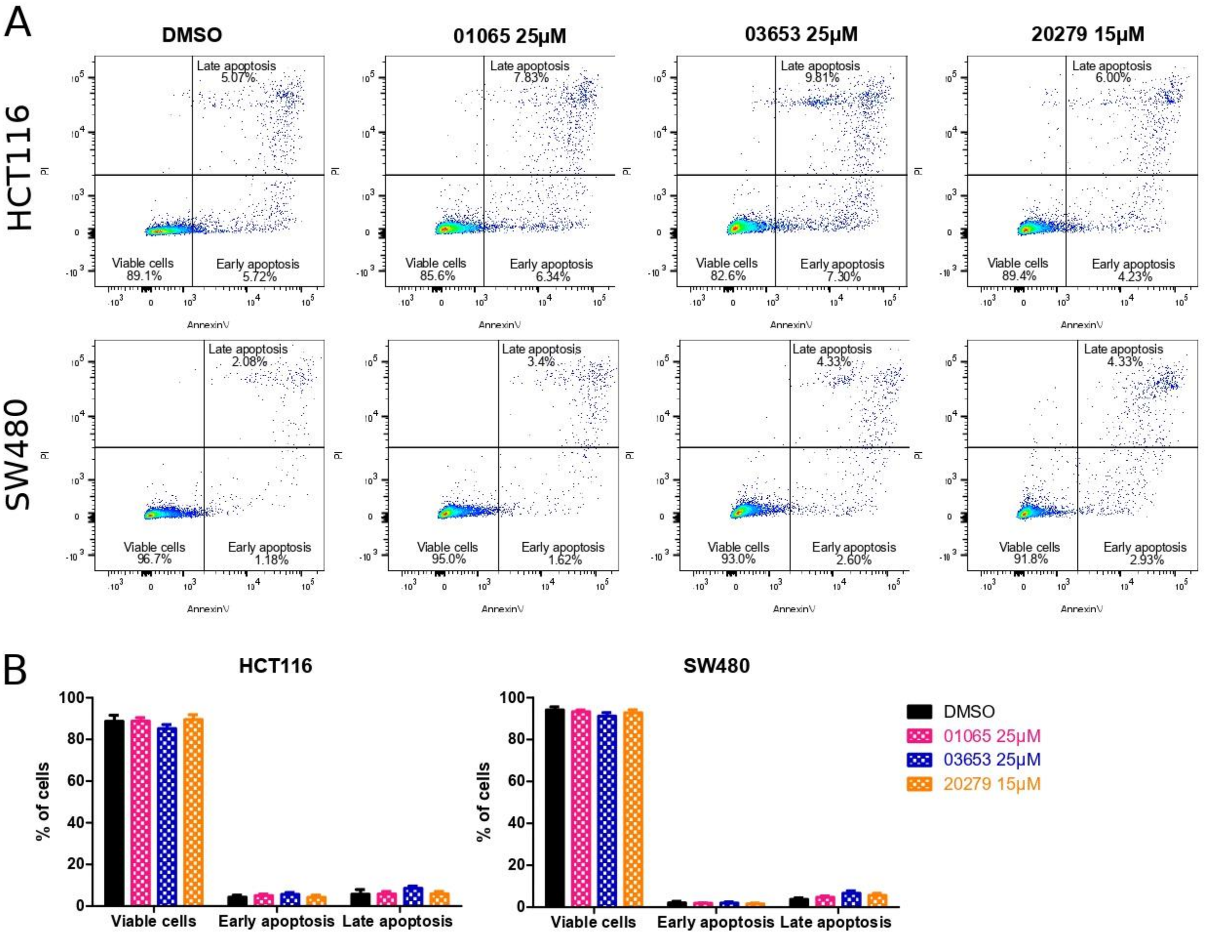

**Figure S5. PTK7/β-catenin inhibitors do not induce cell apoptosis following cell cycle arrest.** (A-B) Effect of compounds 01065, 03653 and 20279 on apoptosis of HCT116 and SW480 cells. Cells were treated for 48h with the different compounds and stained with Annexin V-FITC and propidium iodide before being quantified by flow cytometry. (A) Dot plots were divided into four quadrants to indicate viable cells, early apoptosis and late apoptosis. (B) Cell viability distribution is represented as histogram of the percentage of viable, early apoptotic and late apoptotic cells. Results are expressed as mean of 3 independent experiments ± SEM and are not statistically different between the DMSO and treated groups.

#### III. Supplementary Tables

**Table S1: Small molecule inhibitors targeting β-catenin/TCF4 interaction tested in NanoBRET™ for their cross-reactivity on the PTK7/β-catenin interaction.**

| Compound name | Structure | Specificity | IC50 (µM) NanoBRET PTK7/β-catenin | References |
|---|---|---|---|---|
| **PKF118-310 (Toxoflavin)** | 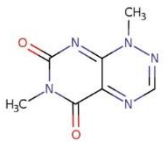 | TCF4 | 101.5 | 8 |
| **LF3** | 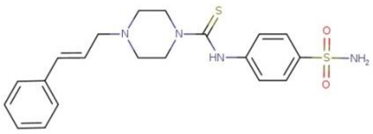 | TCF4<br>Lef1 | 111.7 | 9 |
| **iCRT3** | 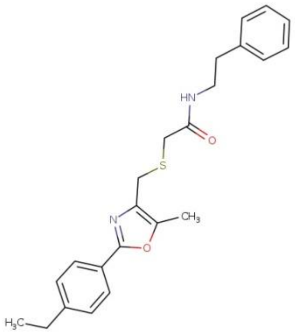 | TCF4<br>Ecadherin +/-<br>αcatenin +/- | 64.19 | 10 |
| **iCRT14** | 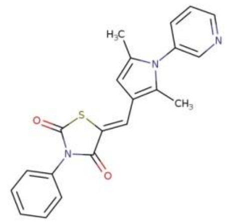 | TCF4<br>Ecadherin +/-<br>αcatenin +/- | 101.1 | 10 |
| **R999636** | 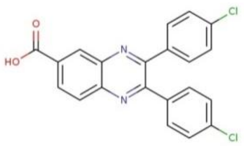 | TCF4<br>E-cadherin<br>APC +/- | 89.48 | 11 |
| **L338192** | 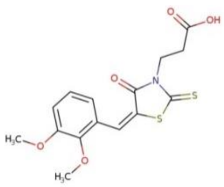 | TCF4<br>APC<br>E-cadherin +/- | N/D | 11 |

| | | | | |
|---|---|---|---|---|
| **PNU-47654** | 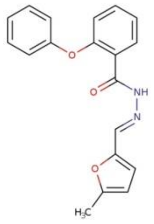 | No data | N/D | 12 |

**Table S2: Selection of compounds from *Vitas-M Laboratory, Ltd.* to probe the structure-activity relationship (SAR)**

| Compound name | Hits | Structure | Molecular Weight | IC50 (µM) NanoBRET |
|---|---|---|---|---|
| **20238** | 01065 | 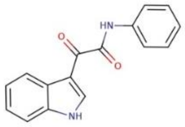 | 264.28 | 177.1 |
| **20269** | 01065 | 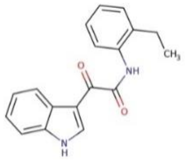 | 292.34 | **19.39*** |
| **20270** | 01065 | 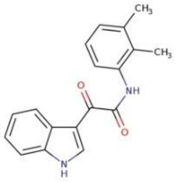 | 292.34 | N/D |
| **20271** | 01065 | 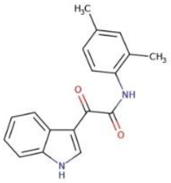 | 292.34 | N/D |
| **20272** | 01065 | 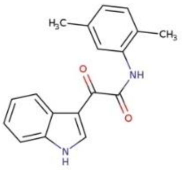 | 292.34 | N/D |
| **20273** | 01065 | 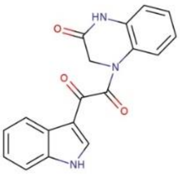 | 319.32 | $2.6X10^{9}$ |
| **20274** | 01065 | 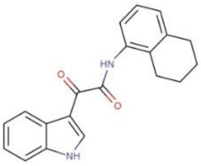 | 318.38 | **12.66*** |
| **20285** | 1065 | 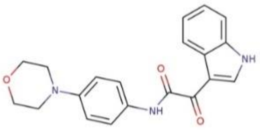 | 349.39 | N/D |

| | | | | |
|---|---|---|---|---|
| **20240** | 02161 | 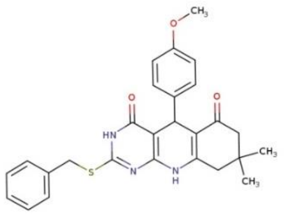 | 473.59 | 1x$10^5$ |
| **20241** | 02161 | 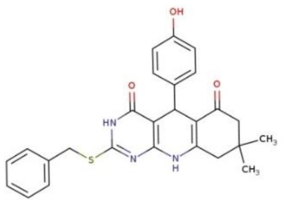 | 459.56 | 142.3 |
| **20242** | 02161 | 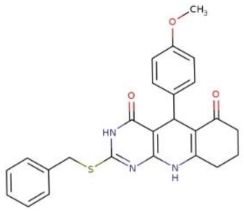 | 445.54 | 94.02 |
| **20243** | 02161 | 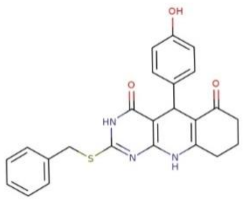 | 431.51 | 3.5x$10^4$ |
| **20244** | 02161 | 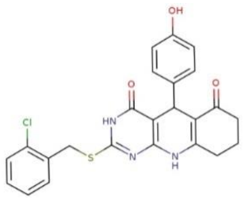 | 465.95 | 174.2 |
| **20245** | 02161 | 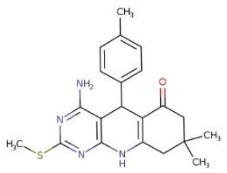 | 380.51 | 155.1 |
| **20246** | 02161 | 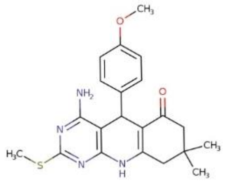 | 396.51 | N/D |
| **20247** | 02161 | 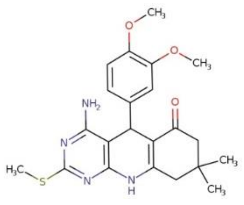 | 426.54 | N/D |

| | | | | |
|---|---|---|---|---|
| **20248** | 02161 | 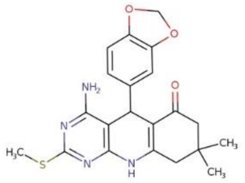 | 410.49 | 117.6 |
| **20249** | 02161 | 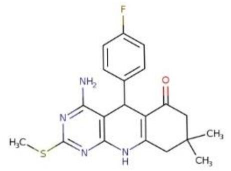 | 384.47 | 137.5 |
| **20250** | 02161 | 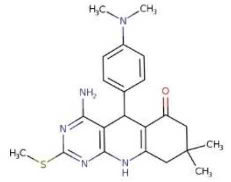 | 409.55 | 252.6 |
| **20251** | 02161 | 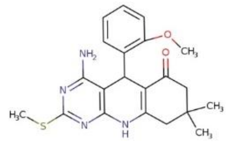 | 396.51 | $1.5x10^4$ |
| **20252** | 02161 | 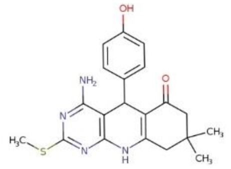 | 382.48 | N/D |
| **20254** | 03653 | 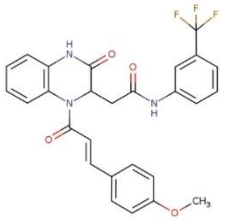 | 509.48 | 86.89 |
| **20255** | 03653 | 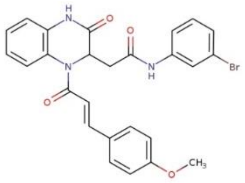 | 520.38 | N/D |
| **20256** | 03653 | 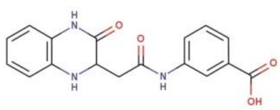 | 325.32 | N/D |
| **20257** | 03653 | 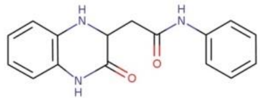 | 281.32 | N/D |

| | | | | |
|---|---|---|---|---|
| **20262** | 03653 | 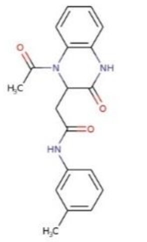 | 337.38 | N/D |
| **20263** | 03653 | 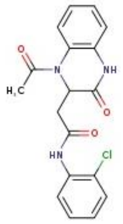 | 357.79 | 107.7 |
| **20275** | 03653 | 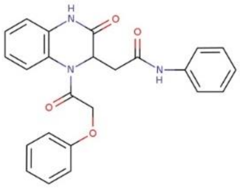 | 415.45 | 9x10$^5$ |
| **20277** | 03653 | 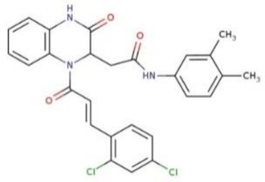 | 508.4 | 49.38 |
| **20278** | 03653 | 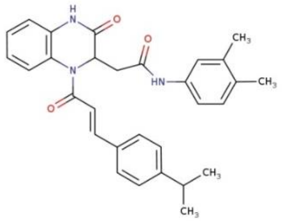 | 481.6 | **37.88*** |
| **20279** | 03653 | 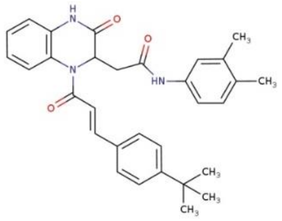 | 495.62 | **17.28*** |
| **20280** | 03653 | 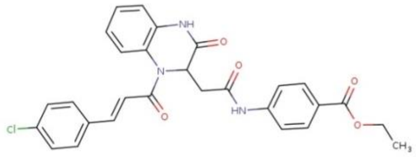 | 517.97 | 97.42 |
| **20239** | 04967 | 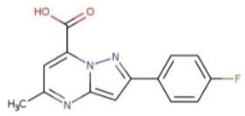 | 271.25 | 2204 |

| | | | | |
|---|---|---|---|---|
| **20266** | 04967 | 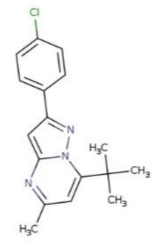 | 299.8 | $1x10^7$ |
| **20276** | 04967 | 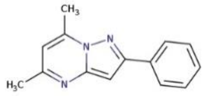 | 223.28 | $1x10^4$ |
| **20281** | 04967 | 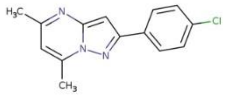 | 257.72 | $1.2x10^5$ |
| **20284** | 04967 | 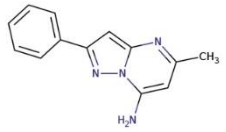 | 224.27 | $4.5x10^5$ |
| **20260** | 05720 | 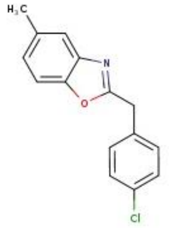 | 257.72 | 344.9 |
| **20268** | 05720 | 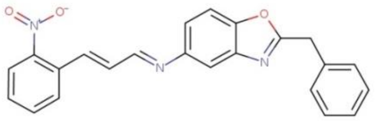 | 383.41 | 57.31 |
| **20282** | 05720 | 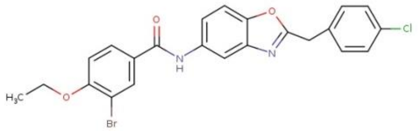 | 485.76 | 51.13 |
| **20283** | 05720 | 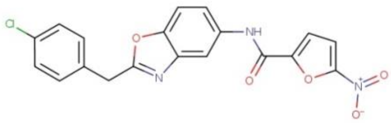 | 397.77 | 29.3 |
| **20286** | 05720 | 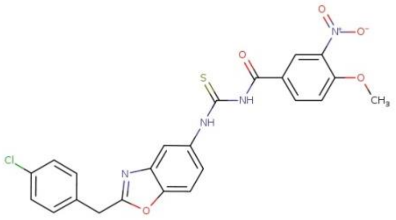 | 496.92 | 105.7 |

| | | | | |
|---|---|---|---|---|
| **20287** | 05720 | 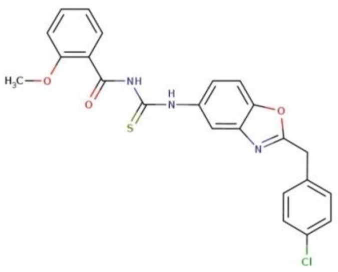 | 451.93 | N/D |
| **20258** | 06062 | 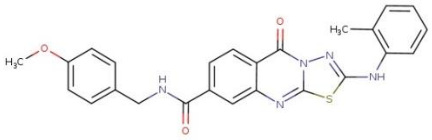 | 471.54 | 55.52 |
| **20259** | 06062 | 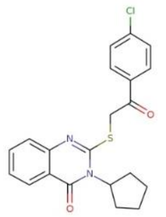 | 398.91 | $6.4\text{x}10^{24}$ |
| **20261** | 06062 | 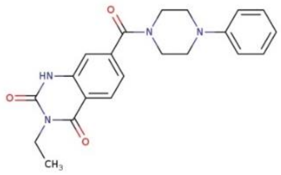 | 378.43 | N/D |
| **20264** | 06062 | 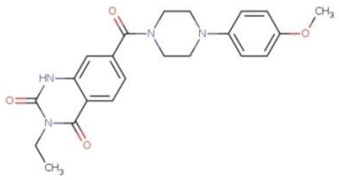 | 408.46 | 125 |
| **20265** | 06062 | 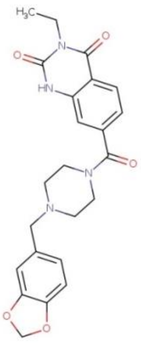 | 436.47 | 86.98 |
| **20267** | 06062 | 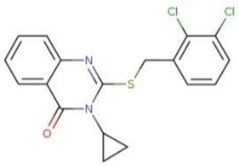 | 377.28 | 129.3 |

| | | | | |
|---|---|---|---|---|
| **20288** | 07912 | 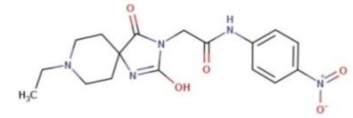 | 375.38 | N/D |
| **20289** | 07912 | 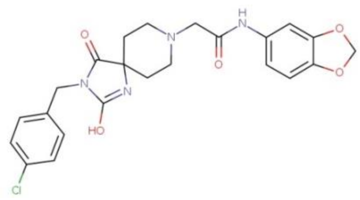 | 470.91 | 78.62 |
| **20290** | 07912 | 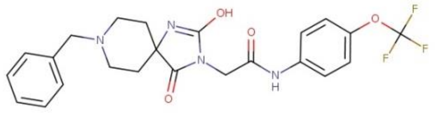 | 476.46 | 77.3 |
| **20291** | 07912 | 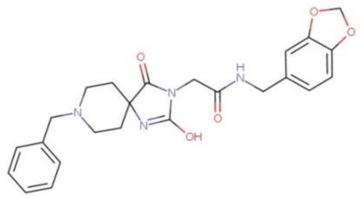 | 450.5 | 757.5 |
| **20292** | 07912 | 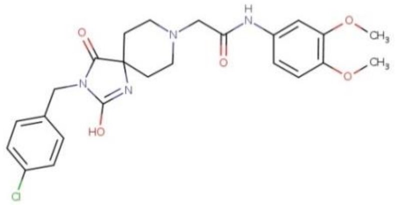 | 486.95 | 175.1 |
| **20293** | 07912 | 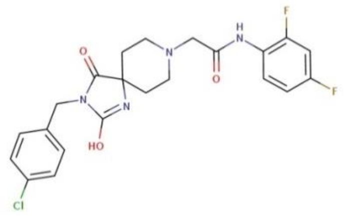 | 462.88 | 75.24 |

**Table S3: List of the 84 key genes responsive to WNT signal transduction in the $RT^2$ Profiler PCR Array and associated functions**

| | |
|---|---|
| **Cell Adhesion Molecules** | ABCB1, CD44, CDH1, CCN2, NRCAM |
| **Cell Migration** | CD44, EFNB1, EGFR, FN1, IRS1, NRCAM, NRP1, PDGFRA, VEGFA |
| **Cell Cycle** | AHR, BIRC5, CCND1, CCND2, EGFR, FOSL1, ID2, MYC, PTGS2 |
| **Differentiation & Development** | Growth Factors : ANGPTL4, BMP4, CCN2, FGF20, FGF4, FGF7, FGF9, GDF5, GDNF, IGF1, IGF2, IL6, JAG1, TGFB3, VEGFA<br>Transcription Factors : CDKN2A, EGR1, NANOG, POU5F1, RUNX2, SIX1, SOX2, SOX9, TBXT, TWIST1<br>Others : ANTXR1, CDON, DAB2, DLK1, EFNB1, FN1, GJA1, ID2, MET, NTRK2, PDGFRA |
| **Calcium Binding & Signaling** | BGLAP, CACNA2D3, CUBN, EGFR |
| **Proteases** | DPP10, MMP3, MMP7, MMP9, PLAUR |
| **Signal Transduction** | WNT signaling : AXIN2, BTRC, CCN4, CCN5, DKK1, FZD7, LRP1, PLPP3, SFRP2, TLE1, WNT3A, WNT5A, WNT9A<br>TGFβ Signaling : BMP4, FST, GDF5, GDNF, TGFB3<br>Hedgehog Signaling : PTCH1, SMO<br>Notch Signaling : JAG1 |
| **Transcription Factors** | WNT Signaling : LEF1, TCF7, TCF7L1, TCF7L2<br>Others : CEBPD, ETS2, KLF5, MYC, PITX2, PPARD, TCF4 |

**Table S4: Primer sequences**

| Gene | | Sequence |
|---|---|---|
| **PTK7** | | For 5'-CAGTTCCTGAGGATTTCCAAGAG-3'<br>Rev 5'-TGCATAGGGCCACCTTC-3' |
| **CTNNB1** | | For 5’-ATTGGCAATGAGCGGTTCCG-3’<br>Rev 5’-AGGGCAGTGATCTCCTTCTG-3’ |
| **AXIN2** | | For 5'-AGTGTGAGGTCCACGGAAAC-3'<br>Rev 5'-CTTCACACTGCGATGCATTT-3' |
| **PLAUR** | | For 5'-CTGCAAGGGGAACAGCAC-3'<br>Rev 5'-GCTTTGGTTTTTCGGTTCG-3' |
| **FOSL1** | | For 5'-CTTGTGAACAGATCAGCCCG-3'<br>Rev 5'-TCCAGTTTGTCAGTCTCCGC-3' |
| **DKK1** | | For 5’-CAACGCTATCAAGAACCTGC-3’<br>Rev 5’-GATCTTGGACCAGAAGTGTC-3’ |
| **NRP1** | | For 5'-AAAACGGTGCCATCCCT-3'<br>Rev 5'-AAGAAGCAGAGTGGGTCGTT-3' |
| **BIRC5** | | For 5'-TGAGAACGAGCCAGACTTGG-3'<br>Rev 5'-TGTTCCTCTATGGGGTCGTCA-3' |
| **KLF5** | | For 5′-GATCTAGATATGCCCAGTTC-3′<br>Rev 5′-CAGCCTTCCCAGGTACACTTG-3′ |
| **MMP7** | | For 5′-GAGTGAGCTACAGTGGGAACA-3′<br>Rev 5′-CTATGACGCGGGAGTTTAACAT-3′ |
| **JAG1** | | For 5'-CAAACACCAGCAGAAAGCCC-3'<br>Rev 5'-TAAGTCAGCAACGGCCTCAG-3' |
| **PTCH1** | | For 5'-ACAACAGGCAGTGGAAATTGGAAC-3'<br>Rev 5'-CCCAGAAGCAGTCCAAAGGTGTAA-3' |
| **Housekeeping genes** | ACTB | For 5’- CCACCGCGAGAAGATGA-3'<br>Rev 5'-CCAGAGGCGTACAGGGATAG-3' |
| | B2M | For 5'-GTCTTTCAGCAAGGACTGGTC-3'<br>Rev 5'-CAAATGCGGCATCTTCAAACC-3' |

**Table S5: Antibodies**

| Target | Clone | Species | Dilution | Cat No. | Company |
|---|---|---|---|---|---|
| **PTK7** | 31G9 | Rat | 1:2000 | In house | |
| **βcatenin** | E-5 | Ms | 1:1000 | Sc-7963 | Santa Cruz |
| **CyclinA2** | BF683 | mS | 1:2000 | CST4656 | Cell Signaling Technology |
| **CyclinD1** | 92G2 | Rb | 1:1000 | CST2978 | Cell Signaling Technology |
| **CyclinE2** | Polyclonal | Rb | 1:1000 | CST4132 | Cell Signaling Technology |
| **CDK2** | 78B2 | Rb | 1:1000 | CST2546 | Cell Signaling Technology |
| **CDK4** | D9G3E | Rb | 1:1000 | CST12790 | Cell Signaling Technology |
| **CDK6** | DCS83 | Ms | 1:2000 | CST3136 | Cell Signaling Technology |
| **p21** | 12D1 | Rb | 1:1000 | CST2947 | Cell Signaling Technology |
| **P27** | D69C12 | Rb | 1:1000 | CST3686 | Cell Signaling Technology |
| **HSP70** | W27 | Ms | 1:1000 | SC-24 | Santa Cruz |
| **Tubulin** | B-5-1-2 | Ms | 1:1000 | T6074 | Sigma Aldrich |

(12) Trosset, J.-Y.; Dalvit, C.; Knapp, S.; Fasolini, M.; Veronesi, M.; Mantegani, S.; Gianellini, L. M.; Catana, C.; Sundström, M.; Stouten, P. F. W.; Moll, J. K. Inhibition of Protein-Protein Interactions: The

Discovery of Druglike Beta-Catenin Inhibitors by Combining Virtual and Biophysical Screening. *Proteins* **2006**, *64* (1), 60–67. https://doi.org/10.1002/prot.20955.
